# The NHSL1-A complex interacts with the Arp2/3 complex and controls cell migration efficiency and chemotaxis

**DOI:** 10.1101/2025.03.13.643034

**Authors:** Shamsinar Jalal, Tommy Pallett, Sheng-yuan Wu, Sreeja B. Asokan, James E. Bear, Matthias Krause

## Abstract

Cell migration is crucial for development and deregulation causes diseases. The Scar/WAVE complex promotes mesenchymal cell migration through Arp2/3 mediated lamellipodia protrusion. We previously discovered that all isoforms of Nance-Horan Syndrome-like 1 (NHSL1) protein interact directly with the Scar/WAVE complex and the NHSL1-F1 isoform negatively regulates Scar/WAVE-Arp2/3 activity thereby inhibiting 2D random cell migration. Here, we investigate the NHSL1-A1 isoform, which contains a Scar homology domain (SHD). The SHD in Scar/WAVE mediates the formation of the Scar/WAVE complex. We found that the SHD of NHLS1-A is sufficient for the formation of an NHSL1-A complex composed of the same proteins as the Scar/WAVE complex, but NHSL1-A replaces Scar/WAVE. NHSL1-A SHD recruits the NHSL1-A complex to lamellipodia, where also the Scar/WAVE complex resides. Scar/WAVE contains a WCA domain, which is phosphorylated by CK2 and recruits and activates the Arp2/3 complex to nucleate branched actin networks supporting lamellipodial protrusion. We identified a WCA domain in NHSL1 which interacts with the Arp2/3 complex. The NHSL1 WCA domain is phosphorylated by GSK3, and this increases the interaction with the Arp2/3 complex. In contrast to NHSL1-F1, the NHSL1-A complex promotes cell migration speed but not cell persistence via the Scar/WAVE complex and potentially via its WCA domain. In addition, the NHSL1-A complex is required for chemotaxis. Mechanistically, the NHSL1-A complex may increase lamellipodial Arp2/3 activity and lamellipodial speed while reducing lamellipodial persistence. Our findings reveal an additional layer of Arp2/3 complex control essential for mesenchymal cell migration highly relevant for development and disease.

## Introduction

Mesenchymal cell migration is driven by the polymerisation of dense, highly branched actin networks that form membrane protrusions, known as lamellipodia, at the leading edge of cells. The Arp2/3 complex, a branched actin nucleator required for formation of these networks, is activated by the C-terminal WH2-Central-Acid (WCA) domain (also known as the VCA domain) of nucleation promoting factors which include N-WASP, WASP, and Scar/WAVE1-3^1–7^. The WH2 domain mediates G-actin recruitment, whilst the central and acidic regions bind to the Arp2/3 complex; the central region forms an amphipathic helix which is required for Arp2/3 complex activation^8,9^.

Activation is regulated by serine phosphorylation. For instance, CK2-mediated phosphorylation of S483/484 in the WCA domain of haematopoietic-specific WASP increases its affinity for the Arp2/3 complex seven-fold^10^. These residues are conserved in N-WASP, and five similarly positioned and CK2-phosphorylated serine residues are found in Scar/WAVE1-3. This regulation, however, is complicated by the fact that phosphorylation of just the first two of these residues (S482/484) conversely inhibits the Arp2/3 complex^11^. In agreement, serine residues in the acidic region of the WCA of *Dictyostelium* Scar are basally phosphorylated and non-phosphorylatable Scar mutants caused large, persistent lamellipodia and persistent cell migration^12^.

Scar/WAVE1-3 form a heteropentameric Scar/WAVE complex (also known as the WAVE regulatory complex, or WRC) along with CYFIP1/2, Nap1, Abi1-3, and HSPC300. The WCA domain is not accessible in the autoinhibited conformation of the Scar/WAVE complex, but this inhibition is relieved by active-Rac binding to CYFIP1/2, tyrosine phosphorylation, and PIP3 binding to Scar/WAVE1-3^13–16^. Once activated, the Scar/WAVE complex drives the complete activation of the Arp2/3 complex and thereby promotes mesenchymal cell migration^1–5,17^.

Cell migration is essential for embryonic development, and hence its dysregulation contributes to developmental defects. Nance-Horan Syndrome is a rare X-linked developmental disorder^18^ caused by mutations in the *NHS* gene. Affected individuals display facial and dental abnormalities, cataracts, and cognitive impairment^19,20^. Nance-Horan Syndrome-like 1 (NHSL1) protein belongs to the poorly characterised Nance-Horan syndrome protein family together with the Nance-Horan Syndrome (NHS), NHSL2^21^, and NHSL3 (HGNC ID:29301)^22^ proteins. NHSL1 has 25 predicted isoforms in GenBank (GeneID: 57224): the result of the alternative splicing of exons 1 and the skipping of exons 3 and/or 6^21,23^. We propose a systematic nomenclature for NHSL1 isoforms: a letter A-H specifying the eight alternatively spliced exon 1 (exon1a-exon1h) and a number, 1-4, indicating (1) the skipping of exons 3 and 6, (2) the skipping of exon 6 only, (3) the skipping of exon 3 only, and (4) the inclusion of both exons 3 and 6 respectively. For example, NHSL1-A1 includes exon 1a, and omits both exon 3 and exon 6 (Fig. S1A).

Exon 1a harbours a Scar Homology domain (SHD), also found in Scar/WAVE proteins (Fig. 1A, S1B,C). In Scar/WAVE proteins the SHD facilitates the formation of a heterotrimer with Abi and HSPC300 which interacts with the CYFIP-Nap1 dimer to form the full Scar/WAVE complex^14^. Consequently, it may be possible for NHSL1 isoforms containing this exon (NHSL1-A isoforms) to also form a trimeric sub-complex with Abi and HSPC300 and thereby form an analogous complex to the Scar/WAVE complex, in which the Scar/WAVE protein is substituted for NHSL1-A.

**Figure 1:**
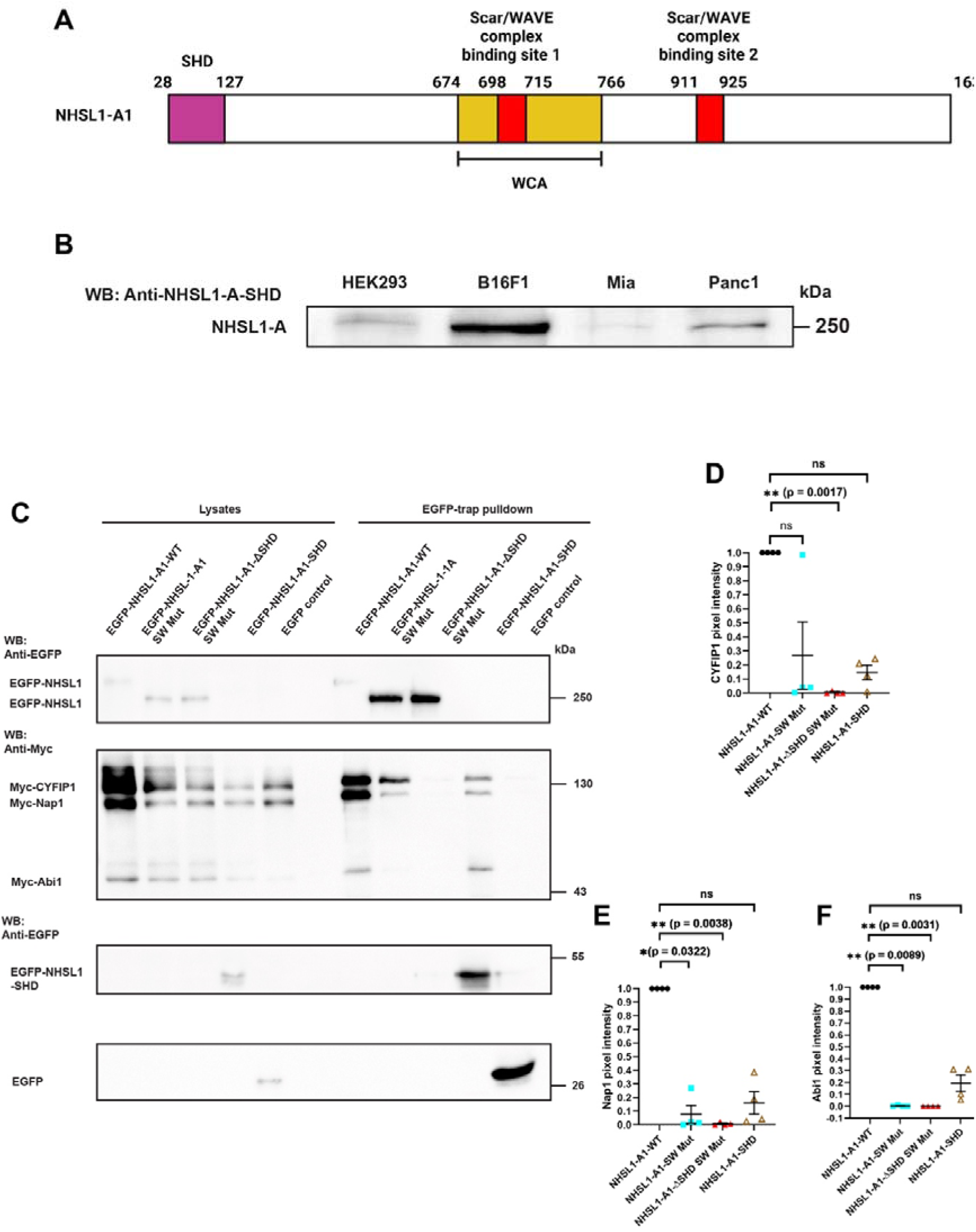
NHSL1-A harbours a Scar homology domain allowing it to form the NHSL1-A complex. **(A)** Domain organisation of the NHSL1-A1 protein isoform. The Scar Homology Domain (SHD), two Scar/WAVE complex (Abi SH3) binding sites and the WH2-central-acidic (WCA) domain are shown. Numbers above the domains indicate the amino acid sequence start and end positions in NHSL1-A1. **(B)** Western blot showing the expression of NHSL1-A, probed with polyclonal NHSL1 SHD specific antibody. Representative blots from three independent biological experiments. **(C)** HEK293 cells were transfected with EGFP-tagged NHSL1-A1, or NHSL1-A1-SW-Mut, or NHSL1-A1ΔSHD-SW-Mut, or the NHSL1-A-SHD (Scar Homology Domain), or EGFP only as negative control and Myc-tagged Scar/WAVE complex components including CYFIP1, Nap1, and Abi1. EGFP-trap pulldown was done to measure the extent of interaction between the different NHSL1-A constructs and the Scar/WAVE complex components. Co-precipitation was detected via a Western blot with Myc antibody. Representative blots from four independent biological experiments. **(D – F)** Quantification of band intensity from chemiluminescence imaging shown in **(C)** normalised to the EGFP negative control. The values are then normalised to NHSL1-A-WT construct, which is set at 1.0. Results are mean values ± SEM (error bars); Kruskal-Wallis test: **(D)** For Myc-tagged CYFIP1, NHSL1-A1-WT vs NHSL1-A1-ΔSHD-SW-Mut: \*\**P* = 0.0017; **(E)** For Myc-tagged Nap1, NHSL1-A1-WT vs NHSL1-A1-SW-Mut: \**P* = 0.0322; NHSL1-A1-WT vs NHSL1-A1-ΔSHD-SW-Mut: \*\**P* = 0.0038; **(F)** For Myc-tagged Abi1, NHSL1-A1-WT vs NHSL1-A1-SW-Mut: \**P* = 0.0089; NHSL1-A1-WT vs NHSL1-A1-ΔSHD-SW-Mut: \*\**P* = 0.0031; *ns* = no significant difference.

**Figure S1:**
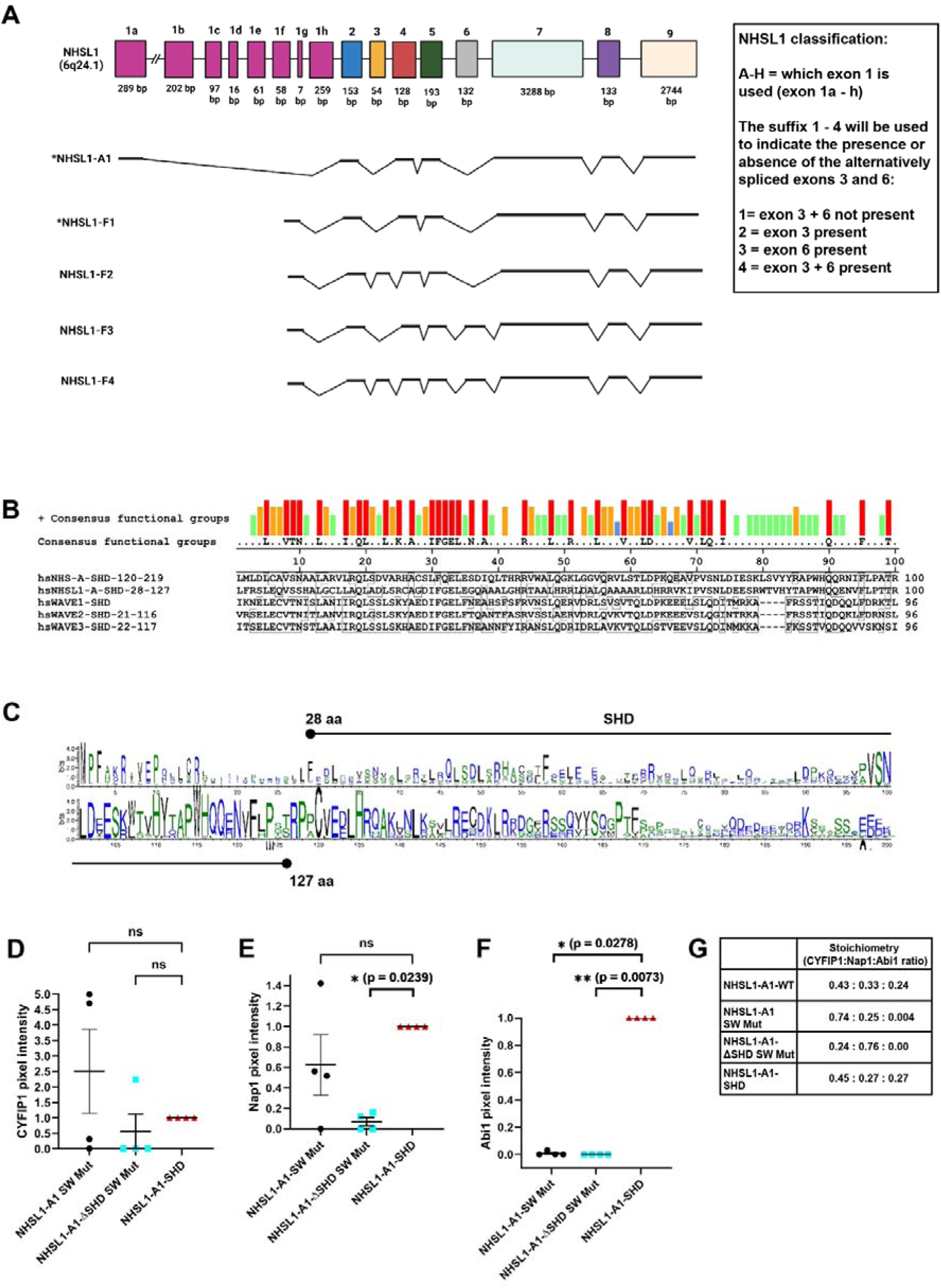
NHSL1 isoform nomenclature, NHSL1-A Scar homology domain alignments, and quantification of NHSL1-A complex interactions. **(A)** Genomic structure of the NHSL1 gene and nomenclature of its isoforms. NHSL1-A1 includes exon 1a, which encodes for most of the SHD. The NHSL1-F1, -F2 and -F4 isoforms all contain exon 1f. The (*) sign indicates the isoforms of NHSL1 being studied in this and our previous study^25^. **(B)** Clustal V sequence alignment of amino acid residues in the SHD of NHS-A, NHSL1-A, and Scar/WAVE1–3. Red indicates conservation between groups of strongly similar properties with Gonnet PAM 250matrix scoring more than 0.5, while blue and green represent conservation between groups of weakly similar properties with Gonnet PAM 250 scoring less than or equal to 0.5. **(C)** Deep MSA conservation of amino acid residues of SHD of NHSL1-A (NCBI reference sequence: XP047275069.1). The SHD of NHSL1-A is from amino acid 28 to amino acid 127. The MSA alignment was done on the Deep MSA platform built by the Zhang lab^55^. Deep MSA is a sensitive multiple sequence alignment open-source program that ‘digs deep’ into the alignments by merging sequences from three whole-genome and meta genome databases through a hybrid homology-detection algorithm. **(D–F)** Quantification of band intensity from chemiluminescence imaging shown in Figure 1(C) normalised to the EGFP negative control. These represent the same data as in Figure 1 (D-F) but with the values normalised to the NHSL1-A1-SHD construct, which is set at 1.0. Results are mean values ± SEM (error bars); Kruskal-Wallis test for all three analyses against NHSL1-A1-SHD. **(G)** Table showing stoichiometry ratio between CYFIP1, Nap1, and Abi1 of each NHSL1-A1 construct from Figure 1(C) and quantified from Figure 1**(D – F)**. Results are from four independent biological repeats.

During our investigation another group reported the existence of an NHSL1-A complex, which they termed the ‘WAVE-shell complex’^24^. Given that this complex does not contain WAVE (Scar/WAVE) we considered this to be misleading and therefore propose the name ‘NHSL1-A complex’. The study by Wang et al. explored the function the PPP2R1A scaffolding unit of the PP2 phosphatase complex and showed that it interacts with the NHSL1-A complex^24^. The authors confirmed our previous finding that knockdown or knockout of all NHSL1 isoforms increased cell migration efficiency^25^ but did not explore the function of the NHSL1-A complex in cell migration – this is the focus of our present study.

We previously showed that all NHSL1 isoforms interact with the Scar/WAVE complex via two Abi SH3 domain binding sites located in exon 6^25^. In our previous study we focussed on the NHSL1-F1 isoform and found that it negatively regulates cell migration via the Scar/WAVE complex, with which it colocalises at the very edge of protruding lamellipodia^25^. Mechanistically, NHSL1-F1 reduced cell migration efficiency by impeding Arp2/3 activity thereby impairing the stability of lamellipodial protrusions^25^.

Here we independently identify the NHSL1-A complex, and show that in contrast to NHSL1-F1, the NHSL1-A complex promotes cell migration speed via the Scar/WAVE complex and is required for chemotaxis. The NHSL1-A complex localises to the edge of lamellipodia, as well as to vesicles, and its recruitment to lamellipodia depends on its SHD but not on the interaction with the Scar/WAVE complex via Abi. This suggests that both the Scar/WAVE complex and the NHSL1-A complex are independently recruited to lamellipodia. Despite this, we surprisingly observe that interfering with the Scar/WAVE complex interaction did result in a shift in NHSL1-A localisation to tips of microspikes and filopodia. We also uncover that all NHSL1 isoforms harbour a WCA domain in a central region of the protein encoded by exon 6. We show that this domain interacts with the Arp2/3 complex, and that this interaction is upregulated via its phosphorylation by the serine/threonine kinase GSK3. This evidence suggests that the NHSL1-A complex may be functionally equivalent to the Scar/WAVE complex.

Re-expression of NHSL1-A1 in pan-isoform NHSL1 CRISPR knockout cells induced dynamically protruding and retracting lamellipodia and “aster-like” F-actin-rich vesicles close to the leading edge. This suggests that the NHSL1-A complex reorganises lamellipodial actin and thereby increases lamellipodial speed while reducing lamellipodial stability.

## Results

### NHSL1-A harbours a Scar homology domain allowing it to form the NHSL1-A complex

To explore the expression of NHSL1-A in cell lines, we generated an antibody against the SHD which is specific to the NHSL1-A isoform. Western blot analysis of B16F1 melanoma cell lysates revealed that this NHSL1 isoform is highly expressed in this cell line and at lower levels also in other cell lines (Fig. 1A,B). B16F1 cells are widely used for studying lamellipodia and mesenchymal cell migration^15,26–28^ and are thus used throughout this study.

To investigate the function of the NHSL1-A complex, we created several mutated NHSL1-A1 cDNAs. We mutated the two known Abi SH3 domain binding sites^25^ in full length NHSL1-A1 which prevents interaction with the entire Scar/WAVE complex (NHSL1-A1-SW-Mut). We created another construct in which we deleted the SHD domain and mutated the two Abi SH3 domain binding sites (NHSL1-A1-DSHD-SW-Mut), which prevents the formation of the NHSL1-A complex, and which does not allow interaction with the Scar/WAVE complex. We also created a construct just with the SHD domain of NHSL1 (NHSL1-A-SHD).

To explore whether a complex harbouring NHSL1-A can form, containing all components of the Scar/WAVE complex except Scar/WAVE itself, we expressed N-terminally EGFP-tagged NHSL1-A1 wild-type (EGFP-NHSL1-A1-WT), the abovementioned mutant constructs, or EGFP only as negative control together with Myc-tagged Scar/WAVE complex components Nap1, CYFIP1, Abi1, and HSPC300 in HEK cells. EGFP-tagged NHSL1-A1 proteins were pulled down from lysates using EGFP-trap beads and interaction with Myc-tagged Scar/WAVE complex components was evaluated by western blot with an anti-Myc antibody. This analysis revealed that full length NHSL1-A1 and NHSL1-A1-SW-Mut interact with components of the Scar/WAVE complex in the absence of Scar/WAVE (Fig. 1C-F). This was no longer the case when the SHD was deleted, and the SW binding site mutated (Fig. 1C-F). This suggests that NHSL1-A1 can form a complex with Nap1, CYFIP1, Abi1, and most likely HSPC300. The SHD of NHSL1-A is sufficient for the formation of the NHSL1-A complex as it also co-precipitates Nap1, CYFIP1, and Abi1 (Fig. 1C-F). Our results are consistent with a recent report also documenting the existence of the NHSL1-A complex^24^.

### All NHSL1 isoforms contain a WCA domain which mediates interaction with the Arp2/3 complex

In the Scar/WAVE complex, the Scar/WAVE protein mediates the function of the complex: its WCA domain recruits the Arp2/3 complex to the leading edge of cells and activates it thereby inducing branched actin nucleation and subsequent lamellipodial protrusion^1^. This raises the question of what mediates the function of the NHSL1-A complex and the intriguing possibility that NHSL1 harbours a WCA domain as well. In all eukaryotic nucleation promoting factors the WCA domain is found at the C-terminal end of the proteins^1,3,7,29^, but this is not the case for NHS family proteins. However, the first identified activator of the Arp2/3 complex, the bacterial protein ActA, harbours a WCA in the N-terminus of the protein^30–33^. Careful analysis of the NHSL1 sequence revealed a putative central WCA (aa690-765). The NHSL1 WCA contains sequence homology to Scar/WAVE, WASP families, and ActA in their respective WH2, central and acidic regions (Fig. 2A, S2A).

**Figure 2:**
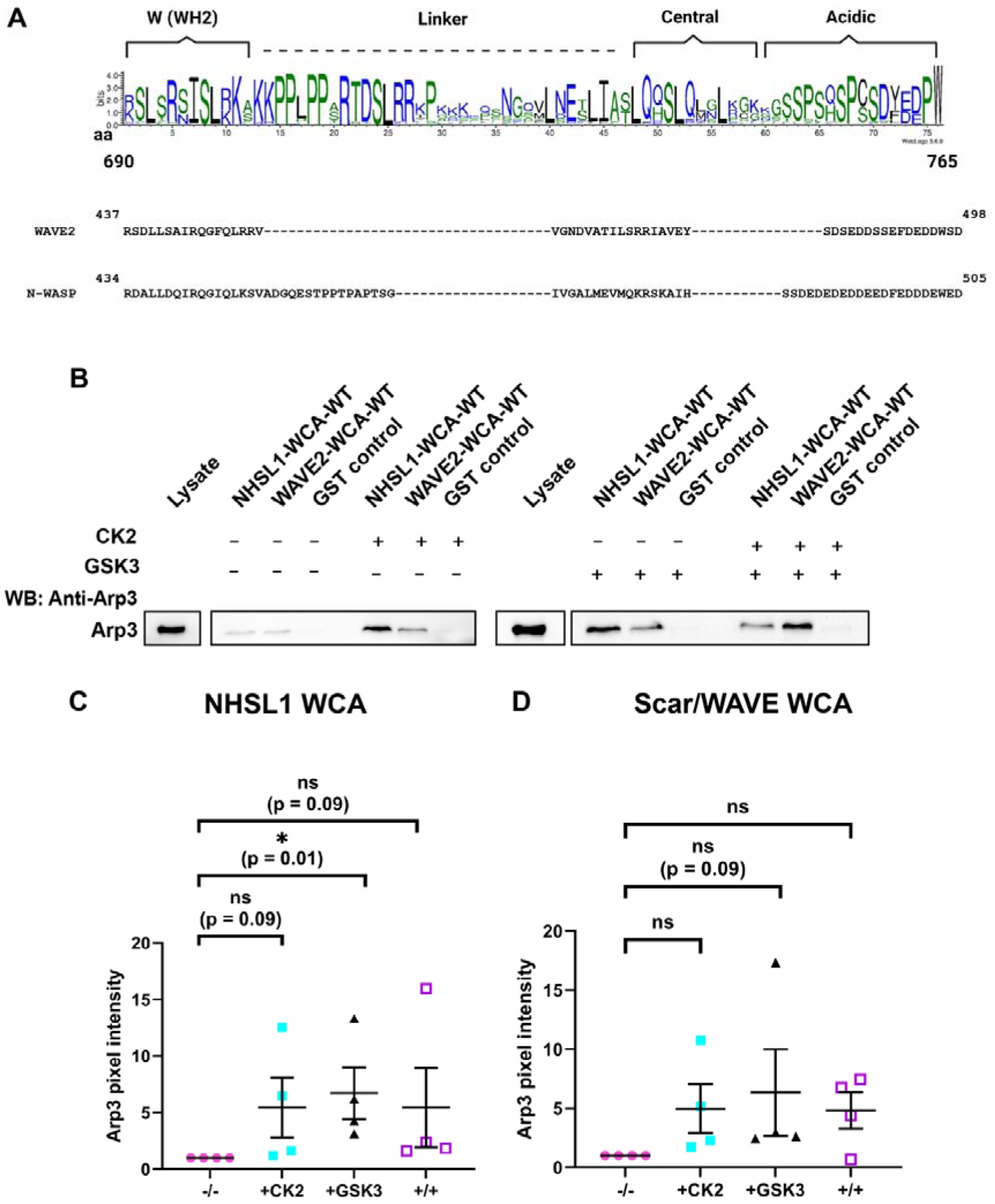
All NHSL1 isoforms contain a WCA domain which mediates interaction with the Arp2/3 complex. **(A)** DeepMSA sequence alignment showing conservation of amino acid residues in the WCA domain of all NHSL1 isoforms. The MSA alignment was done on the DeepMSA platform built by the Zhang lab. The amino acid sequences of WCA of Scar/WAVE2 and N-WASP are also shown as comparison. **(B)** GST-pulldowns using purified GST-tagged proteins of NHSL1-A1-WCA-WT or Scar/WAVE2-WCA or GST alone as a negative control coupled to Glutathione-sepharose beads from B16F1 cell lysates. Before GST-pulldowns were performed, the purified GST-tagged proteins on the sepharose beads were subjected to a dephosphorylation step using calf intestinal phosphatase (CIP) to remove any phosphorylation from the bacterial host, followed by a phosphorylation step using no kinase treatment (-/-), or purified CK2 alone, or GSK3 alone or both CK2 and GSK3 kinases (+/+). Following GST-pulldowns, endogenous Arp3 was detected in a western blot with Arp3 antibody. Representative blots from four independent biological experiments. **(C)** and **(D)** quantification of band intensity from chemiluminescence imaging from **(B)**, normalised to GST control. The values were then normalised to -/-, which was set at 1. Bars indicate mean ± SEM (error bars). Kruskal-Wallis tests: **(C)** GST-NHSL1-WCA: -/- vs + CK2: ns=0.0901; -/- vs + GSK3: **P=0.0134; -/- vs +/+: ns = 0.0901. Kruskal-Wallis tests: **(D)** GST-Scar/WAVE2- WCA -/- vs + CK2: ns = 0.1553; -/- vs + GSK3: **P = 0.0901; -/- vs +/+: ns = 0.1843.

**Figure S2:**
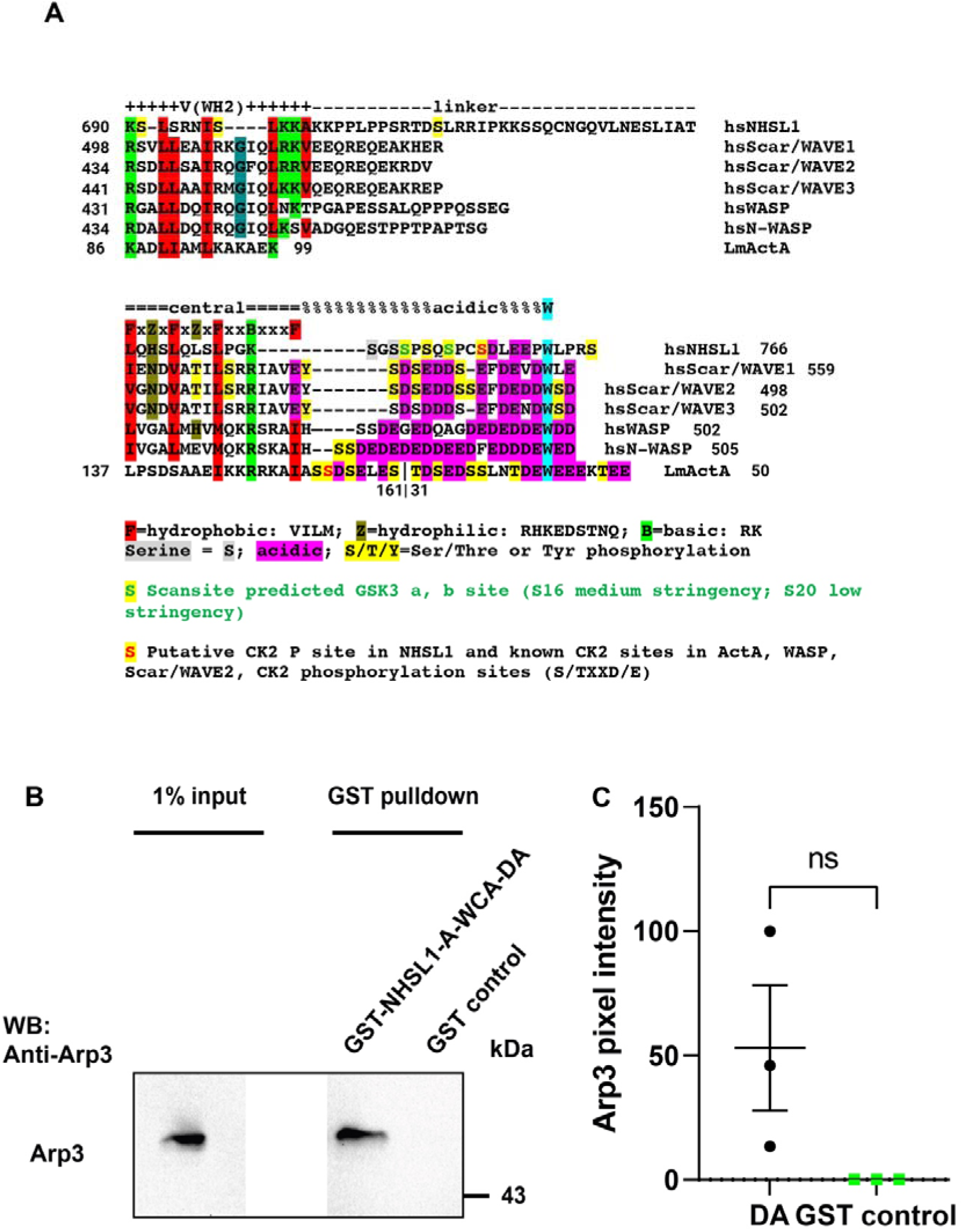
NHSL1 harbours a WCA domain. **(A)** Amino acid sequence alignment of the WCA domains of homo sapiens (hs) NHSL1, Scar/WAVE1–3, WASP, N-WASP and *Listeria monocytogenes* ActA. **(B)** GST-pulldown using purified GST-tagged proteins of dominant active NHSL1-A1-WCA-DA or GST alone as a negative control coupled to Glutathione-sepharose beads from B16F1 cell lysates. Following GST-pulldowns, endogenous Arp3 was detected in a western blot with an Arp3 antibody. The 1% input band and the pulldown bands were taken from the same blot using the same exposure time. Representative blots from three independent biological experiments. **(C)** Quantification of Arp3 band intensity from blot chemiluminescence from **(B)**, normalised to GST control. Results are mean ± SEM (error bars); Unpaired t-test; GST control vs NHSL1-A1-WCA-DA: *ns* = no significant difference. Results are from three independent biological experiments.

The acidic region of NHSL1 WCA harbours a tryptophane, several acidic amino acids, and serine residues in a similar arrangement as in well characterised WCA domains (Fig. 2A, S2A). The latter ones are known to be phosphorylated by the Serine/Threonine kinase CK2 in WASP, Scar/WAVE2, and ActA and this increases the affinity of the WCA for the Arp2/3 complex^10,11,34–36^. Bioinformatics analysis (Scansite, NetPhorest; PhosphositePlus; Kinase library) predicted that CK2 and another Serine/Threonine kinase, GSK3, may phosphorylate the WCA of NHSL1. Mass spectrometry screens revealed that several of the Serine residues in the WCA of NHSL1 are indeed phosphorylated in cells (PhosphositePlus; ^37^) (Fig. S2A).

Thus, we postulated that the WCA of NHSL1 may be regulated by Serine/Threonine phosphorylation to control interaction with the Arp2/3 complex. To test this, we purified the GST-tagged WCA domain of NHSL1, the WCA domain of Scar/WAVE2 as positive control, and GST only as negative control from *E. coli*. These sepharose bead immobilised GST fusion proteins were dephoshophorylated using Calf Intestinal Phosphatase and then *in vitro* phosphorylated with purified CK2, GSK3 or both kinases together. The beads were incubated with B16F1 cell lysates and bound endogenous Arp2/3 complex detected in a western blot with an Arp3 antibody. This analysis revealed that both the NHSL1 WCA domain (Fig. 2B,C) and the Scar/WAVE2 WCA domain (Fig. 2B,D) weakly interact with the Arp2/3 complex without phosphorylation. However, GSK3 phosphorylation resulted in an increased affinity of the NHSL1 WCA domain for the Arp2/3 complex (Fig. 2B,C). Taken together, this suggest that all isoforms of NHSL1 harbour a WCA domain which mediates the interaction with the Arp2/3 complex, and that this interaction is promoted by GSK3 phosphorylation.

To mimic the serine phosphorylation in the NHSL1 WCA, we mutated the serine residues S752E, S756E, and S759E to glutamic acid to create a dominant active NHSL1 WCA (GST-NHSL1-WCA-DA) which may constitutively interact with the Arp2/3 complex. We purified this GST-NHSL1-WCA-DA protein from *E. coli* and tested for interaction with the endogenous Arp2/3 complex from B16F1 cell lysates. The western blots revealed that GST-NHSL1-WCA-DA was able to strongly interact with the Arp2/3 complex (Fig. S2B,C). This provides evidence that the phosphorylation of serine residues in position S752, S756, and S759 promote interaction with the Arp2/3 complex.

### The NHSL1-A complex localises to the very edge of lamellipodia and to vesicles

To explore the function of the NHSL1-A complex in cell migration, we first tested the localisation of our NHSL1-A1 wild-type and mutant constructs in NHSL1 CRISPR knockout B16F1 (NHSL1-KO) cells^25^ to avoid any potential interference with other endogenous NHSL1 isoforms. We co-transfected our EGFP-tagged NHSL1 constructs with LifeAct-mScarlet-I to evaluate any changes in F-actin distribution and dynamics. As a control, we expressed EGFP (Fig. 3A; Supplemental movie 1) which as expected diffusely localises within the cytosol and the nucleus. EGFP-NHSL1-A1-WT localised to the edge of lamellipodia in a similar manner to NHSL1-F1^25^, but also localised strongly to many F-actin-rich vesicles at the leading edge and around the nucleus (Fig. 3B; Supplemental movie 2). In around half of the cells some of the vesicles at the leading edge appear as F-actin rich “asters” from which more vesicles appear to bud off (Fig. S3, arrowhead; Supplemental movie 3) suggesting that these F-actin-rich vesicles may be induced by the expression of wild-type NHSL1-A1.

**Figure 3:**
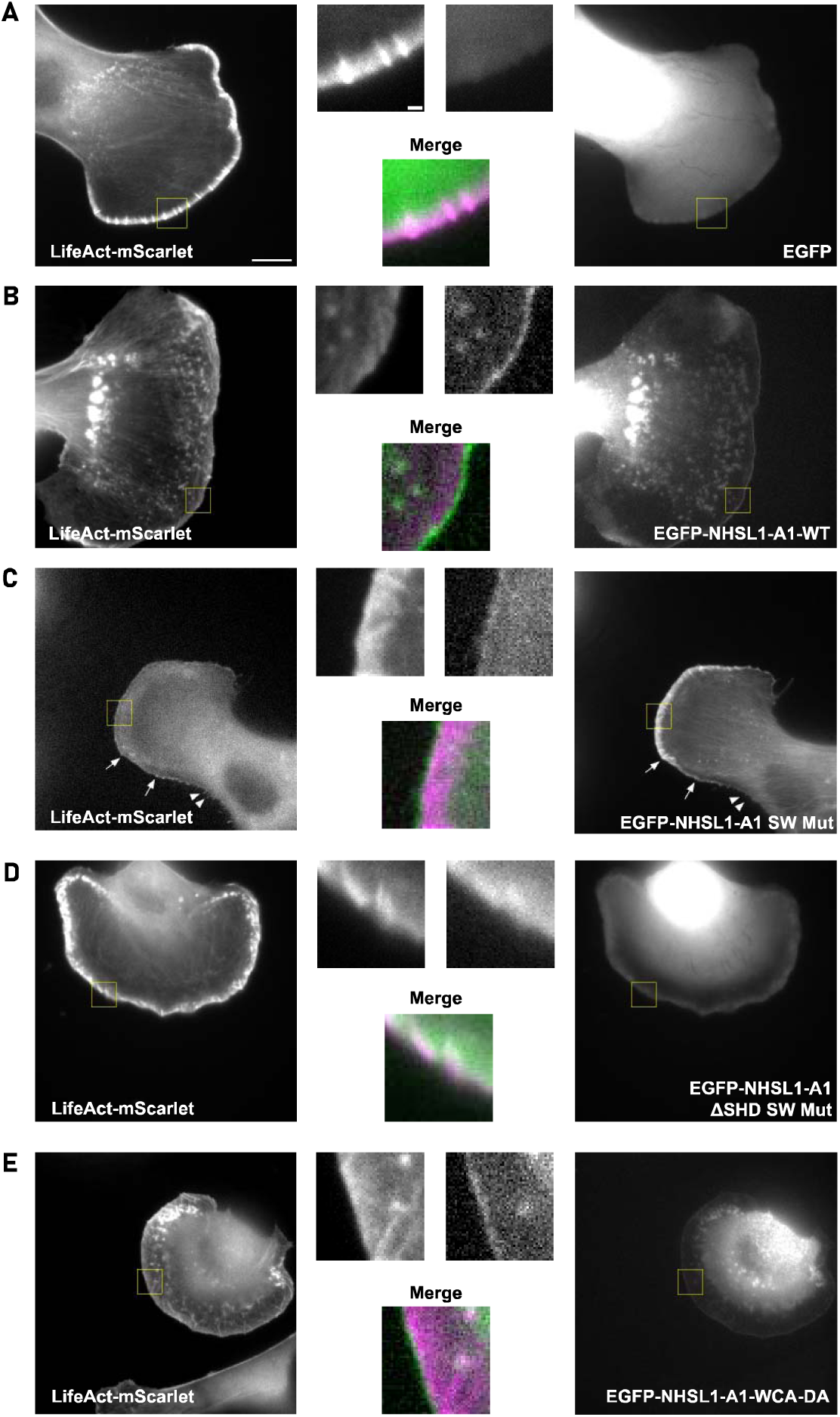
The NHSL1-A complex localises to the edge of lamellipodia and to vesicles. Still images from live-cell imaging showing NHSL1 CRISPR knockout B16F1 cells either expressing EGFP only **(A)**, or **(B – E)** different EGFP-tagged NHSL1-A1 constructs together with LifeAct-mScarlet migrating on laminin. **(B,E)** EGFP-NHSL1-A1-WT and EGFP-NHSL1-A1-WCA-DA localised to the edge of lamellipodia, and vesicles as shown in the insets. **(C)** EGFP-NHSL1-A1-SW-Mut weakly localised to lamellipodia and vesicles but also robustly to tips of microspikes (arrows) and filopodia (arrowheads). **(D)** NHSL1-A1-ΔSHD-SW-Mut localise localised to the cytosol and weakly to vesicles but not to the leading edge. Representative images from four independent biological experiments. Scale bar represents 10 µm. Inset is a magnified view of the white box. Scale bar of inset represents 2 µm.

**Figure S3:**
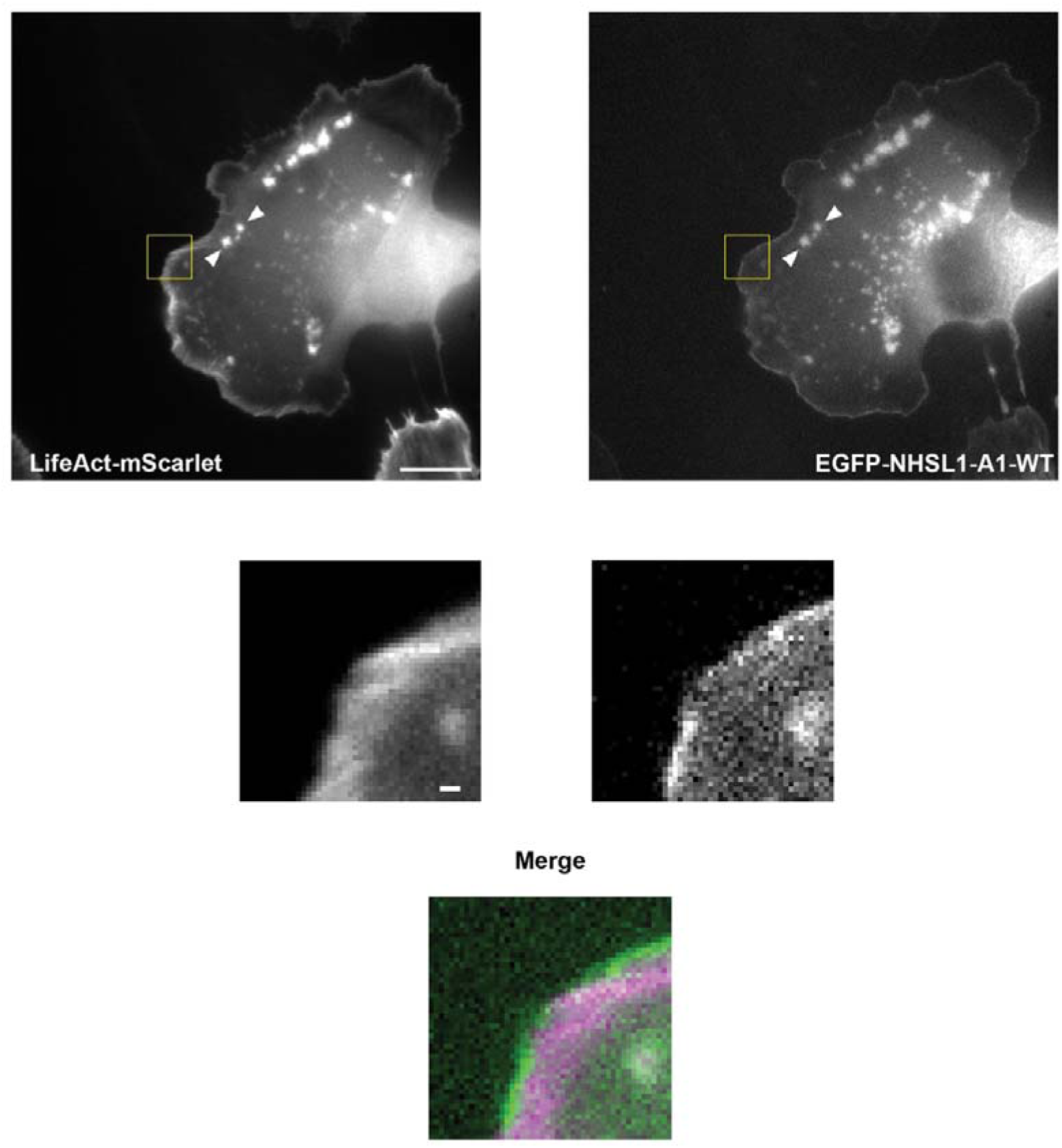
The NHSL1-A complex localises to F-actin-rich “asters” from which vesicles bud off. Still images from live-cell imaging showing NHSL1 CRISPR knockout B16F1 cells expressing EGFP-NHSL1-A1-WT localised to the edge of lamellipodia and in around half of the cells also to F-actin-rich “asters” from which vesicles bud off (arrowheads). Representative images from four independent biological experiments. Scale bar represents 10 µm. Inset is a magnified view of the white box. Scale bar of inset represents 2 µm.

The leading edge of cells transfected with wild-type NHSL1-A1 appeared to be very dynamic in comparison to the NHSL1 knockout cells (Supplemental movies 2 and 3). The EGFP-NHSL1-A1-SW-Mut construct (the variant which cannot bind to the Scar/WAVE complex) localised more weakly to the edge of lamellipodia and to only a few F-actin-rich vesicles compared to wild-type NHSL1-A1. However, in addition NHSL1-A1-SW-Mut localisation was observed at the tips of microspikes (Fig. 3C, arrows) and filopodia (Fig. 3C, arrowheads) (Supplemental movie 4). We did not previously detect the analogous mutant variant of NHSL1-F1 (NHSL1-F1-SW-mut) at microspikes or filopodia^25^ suggesting that the NHSL1-A complex can re-locate to these sites when not bound to the Scar/WAVE complex.

We observed that the EGFP-NHSL1-A1-DSHD-SW-mut (which cannot bind to the Scar/WAVE complex nor form the NHSL1-A complex) did not localise to the very edge of lamellipodia and only weakly to vesicles but instead localised to the cytosol and in cells with high expression also to the nucleus, (Fig. 3D; Supplemental movie 5). The leading edge appeared smoother compared to the more dynamic lamellipodia of the NHSL1-A1-WT expressing cells (Fig. 3B,D; Supplemental movie 5).

Similar to wild-type NHSL1-A1, the WCA dominant active mutant construct (EGFP-NHSL1-A1-WCA-DA) localised to the edge of lamellipodia and to vesicles and appeared to increase the number of F-actin rich vesicles (Fig. 3B,E; Supplemental movie 6) and to induce more dynamic lamellipodia (Supplemental movie 6).

### The NHSL1-A complex promotes cell migration speed via the Scar/WAVE complex

We tested the function of the NHSL1-A complex in random cell migration by re-expressing our NHSL1-A1 wild-type and mutants in the NHSL1 CRISPR knockout B16F1 cells. In this case the proteins were Myc-tagged and expressed from a bicistronic plasmid also encoding the resistance gene for blasticidin. This enabled blasticidin selection for successfully transfected cells to ensure all analysed cells expressed the expected constructs. As control we used the empty Myc-tag-IRES-blasticidin plasmid.

Tracking cell migration behaviour from phase contrast movies revealed that cells re-expressing NHSL1-A1-WT migrated significantly faster than the NHSL1 knockout cells (Fig. 4A; for cell tracks see Fig. S4). This is a surprising result because it is opposite to what we observed for the NHSL1-F1 isoform^25^ and suggests that the NHSL1-A complex functions to promote cell migration. Re-expression of the NHSL1-A-SHD on its own did not increase migration speed indicating that the C-terminus of NHSL1 mediates its function in cell migration (Fig. 4A). In agreement, expression of the NHSL1-A1-SW-Mut did not increase migration speed (Fig. 4A) implying that the increase in cell migration requires the interaction of NHSL1-A with the Scar/WAVE complex. However, the NHSL1-A1-DSHD-SW-Mut increased cell migration speed like NHSL1-A1-WT (Fig. 4A). This was unexpected because this construct does not localise to the edge of lamellipodia (Fig. 3D) and cannot form the NHSL1-A complex (Fig. 1C-F). However, it does localise weakly to vesicles and harbours an intact WCA domain and thus may increase cell migration speed indirectly via Arp2/3 complex mediated vesicle trafficking. Re-expression of NHSL1-A1-WCA-DA did not result in cell speeds significantly different from NHSL1-A1-WT suggesting that the NHSL1 WCA does not have a negative function in cell migration (Fig. 4A). This indicates that the function of the NHSL1-A complex in promoting cell migration speed may be mediated by both its interaction with the Scar/WAVE complex and the Arp2/3 complex.

**Figure 4:**
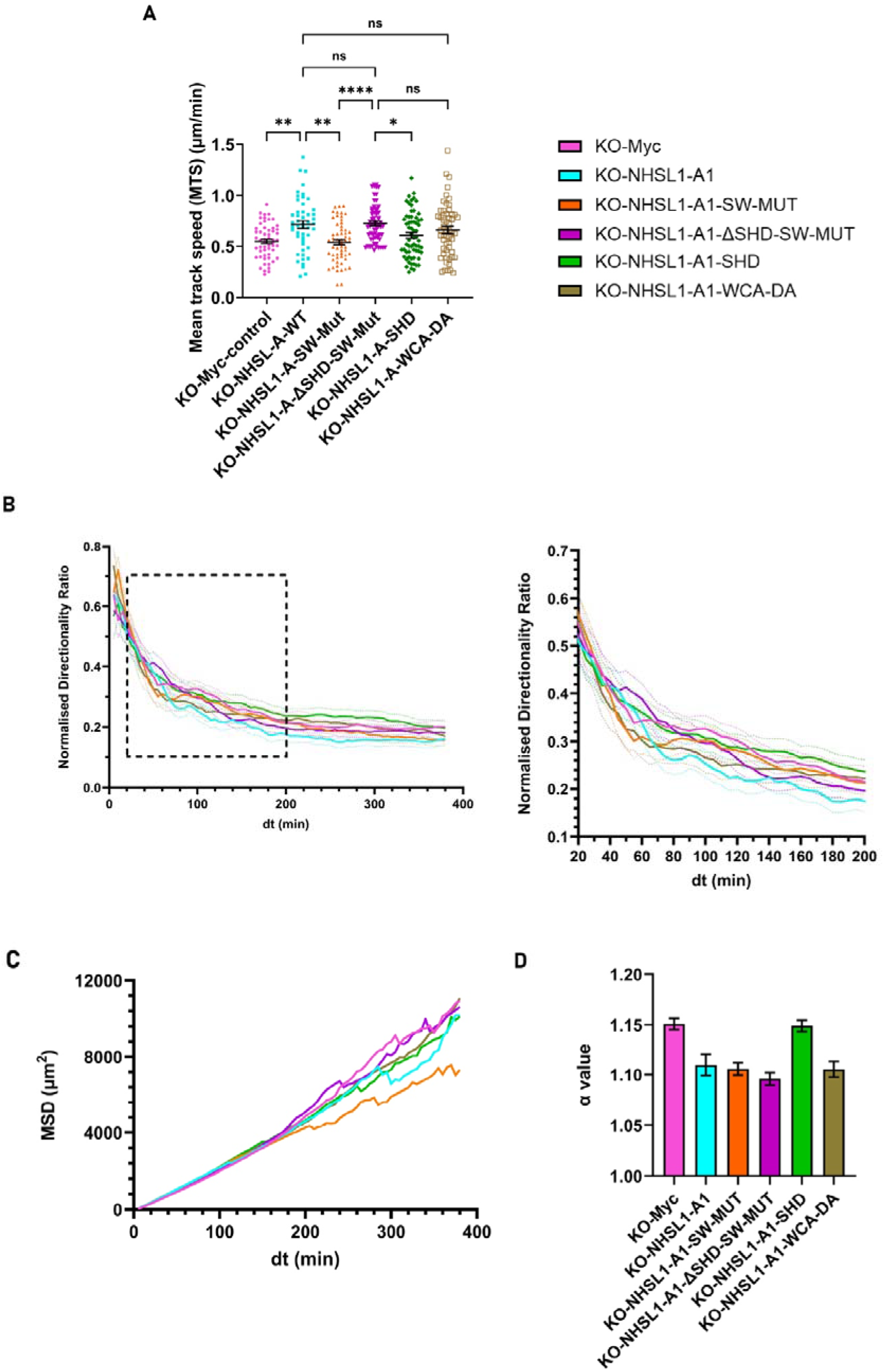
The NHSL1-A complex promotes cell migration speed via the Scar/WAVE complex. **(A)** Mean track speed of randomly migrating NHSL1 CRISPR knockout B16F1 cells expressing Myc alone as control, or NHSL1 CRISPR knockout B16F1 cells expressing either the full-length wild-type NHSL1-A1 or full-length mutant constructs of NHSL1-A1 plated on laminin after selection using a bicistronic blasticidin expression plasmid ensuring all cells analysed expressed NHSL1-A1. Each data point represents the mean speed of a cell. Results are mean ± SEM (error bars). **(B)** The normalised (see methods) directionality ratio was plotted to explore how the cell migration persistence changes over time. Directionality ratio and **(C)** mean square displacement (MSD) were calculated and plotted using excel macros provided in^54^. **(D)** Alpha value (α-value) calculated from the slope of the log-log plot of the MSD of randomly migrating cells. Experiments were performed on *n* = 55 (NHSL1 knockout - Myc only), 49 (NHSL1 knockout - Rescue Myc-NHSL1-A-WT), 58 (NHSL1 knockout - Myc-NHSL1-A-SW-Mut), 62 (NHSL1 knockout - Myc-NHSL1-A-ΔSHD-SW-Mut), 64 (NHSL1 knockout - Myc-NHSL1-A-SHD) and 55 (NHSL1 knockout - Myc-NHSL1-A-WCA-DA) from three independent biological experiments.

**Figure S4:**
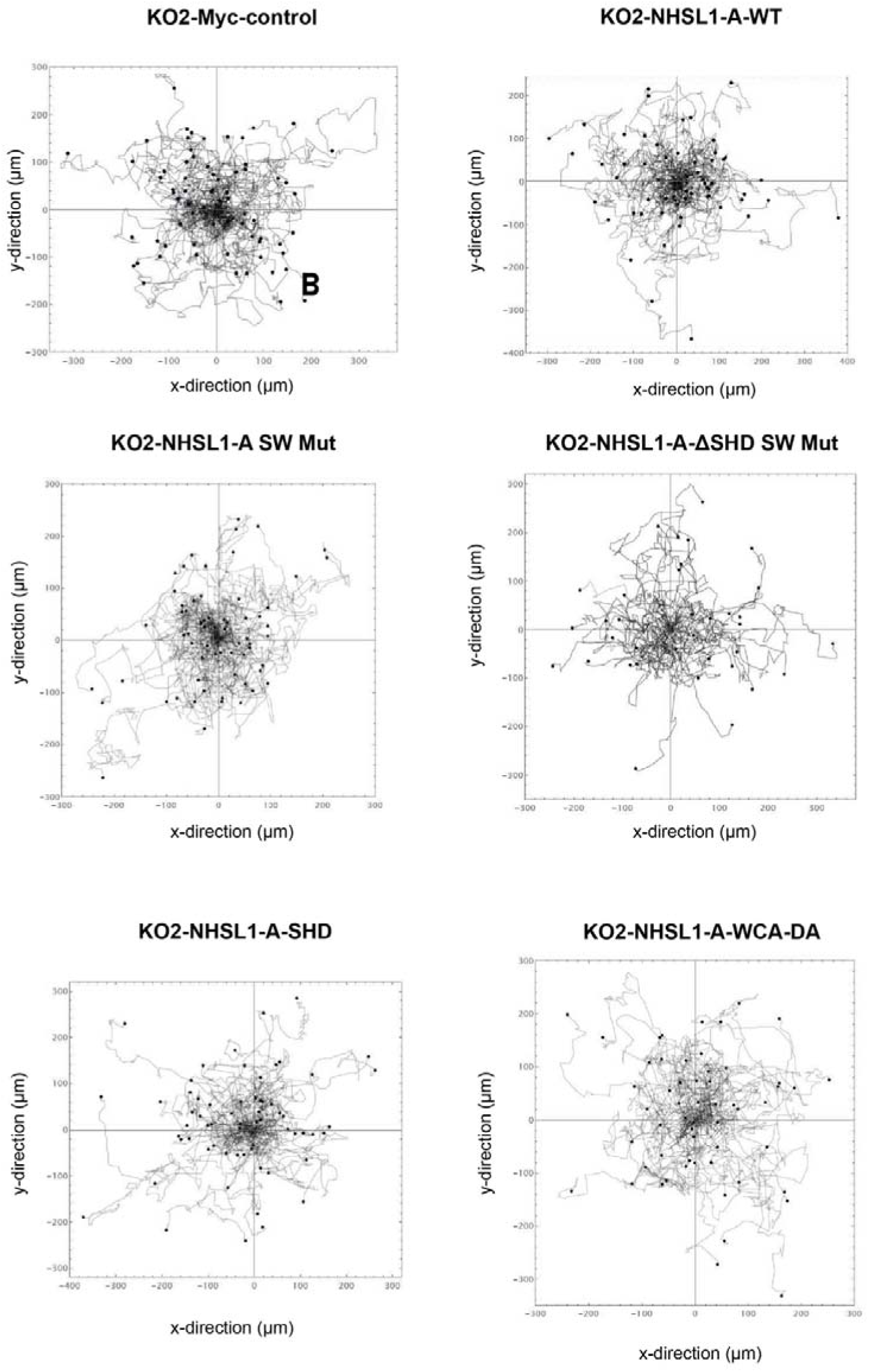
The NHSL1-A complex promotes cell migration speed via the Scar/WAVE complex. Cell tracks of randomly migrating NHSL1 knockout B16F1 cells expressing Myc alone as control, or NHLS1 knockout B16F1 cells expressing either the full-length wild-type NHSL1-A1 or full-length mutant constructs of NHSL1-A1 plated on laminin after selection using a bicistronic blasticidin expression plasmid ensuring all cells analysed expressed NHSL1-A1.

When we analysed the directionality ratio over time, we observed that during early time intervals, which reflect cell migration persistence, the re-expression of NHSL1-A1-WT, NHSL1-A1-SW-Mut, NHSL1-A1-DSHD-SW-Mut, and NHSL1-A1-WCA-DA resulted in the quickest loss (steepest gradient of descent) of directionality compared to the KO-Myc and re-expression of NHSL1-A1-SHD (40<dt<70 min) (Fig. 4B). However, at later intervals (70<dt<200 min) cells re-expressing NHSL1-A1-WT had a reduced directionality, but neither NHSL1-A1-SHD, NHSL1-A1-SW-Mut, nor NHSL1-A1-WCA-DA reduced directionality compared to the NHSL1 knockout cells.

Analysis of mean squared displacement (MSD) which is dependent on speed and persistence only revealed a big reduction in MSD for the re-expression of NHSL1-A1-SW-Mut which reflects its inability to increase migration speed in addition to reducing persistence (short time-interval directionality). Re-expression of NHSL1-A1-WT only moderately reduced MSD which reflects its combination of increased cell migration speed but reduced persistence (Fig. 4C). Estimation of the MSD exponent (alpha-value) from the slopes of the log-log transform of MSD curves characterises the motion of the cells (α = 1 indicates diffusive, or random, motion whilst α = 2 indicates ballistic, or directional, motion)^38,39^. This measure of directionality revealed a similar reduction in cell migration persistence for re-expression of NHSL1-A1-WT, NHSL1-A1-SW-Mut, NHSL1-A1-DSHD-SW-mut, and NHSL1-A1-WCA-DA as observed by measuring directionality over time at shorter interval values. However, re-expression of the NHSL1-A1-SHD did not reduce the alpha value compared to the NHSL1 knockout cells (Fig. 4D).

Taken together, these data suggest that NHSL1-A may inhibit cell migration persistence via other interactors and not via its interaction with the Scar/WAVE and the Arp2/3 complexes (Fig. 4B). There are also clear differences between the regulation of the persistence (defined by shorter time intervals) and longer-term directionality.

### The NHSL1-A complex mediates chemotaxis

It is unknown whether NHSL1 proteins regulate chemotaxis. To explore this, we compared wild-type B16F1 cells with NHSL1 knockout cells both transiently transfected with the empty Myc-tag-IRES-blasticidin vector or the NHSL1 knockout cells re-expressing Myc-NHSL1-A1-WT-IRES-blasticidin. After blasticidin selection, the chemotactic ability of each cell population was analysed in a microfluidic chemotaxis chamber^40^ in a gradient of HGF (Fig. 5A). In the absence of NHSL1 proteins, cell migration speed was significantly reduced from 0.84 ± 0.03 µm/min to 0.67 ± 0.03 µm/min, and re-expression of NHSL1-A1 increased the mean speed back to 0.73 ± 0.04 µm/min µm/min (Fig. 5B), a result which is not statistically significant, but which was in approximate agreement with our results from random migration assays (Fig. 4A). The y-Forward Migration Index (y-FMI) shows that wild-type B16F1 cells chemotax, albeit weakly, up the gradient of HGF with a mean y-FMI of 0.05 ± 0.02. The NHSL1 knockout cells, however, have a mean y-FMI of –0.04 ± 0.02, indicating that they cannot chemotax. This dysfunctional phenotype is rescued significantly by re-expression of wild-type NHSL1-A1 resulting in a mean y-FMI of 0.11 ± 0.03 (Fig. 5C,D) suggesting that the NHSL1-A complex is required for chemotaxis.

**Figure 5:**
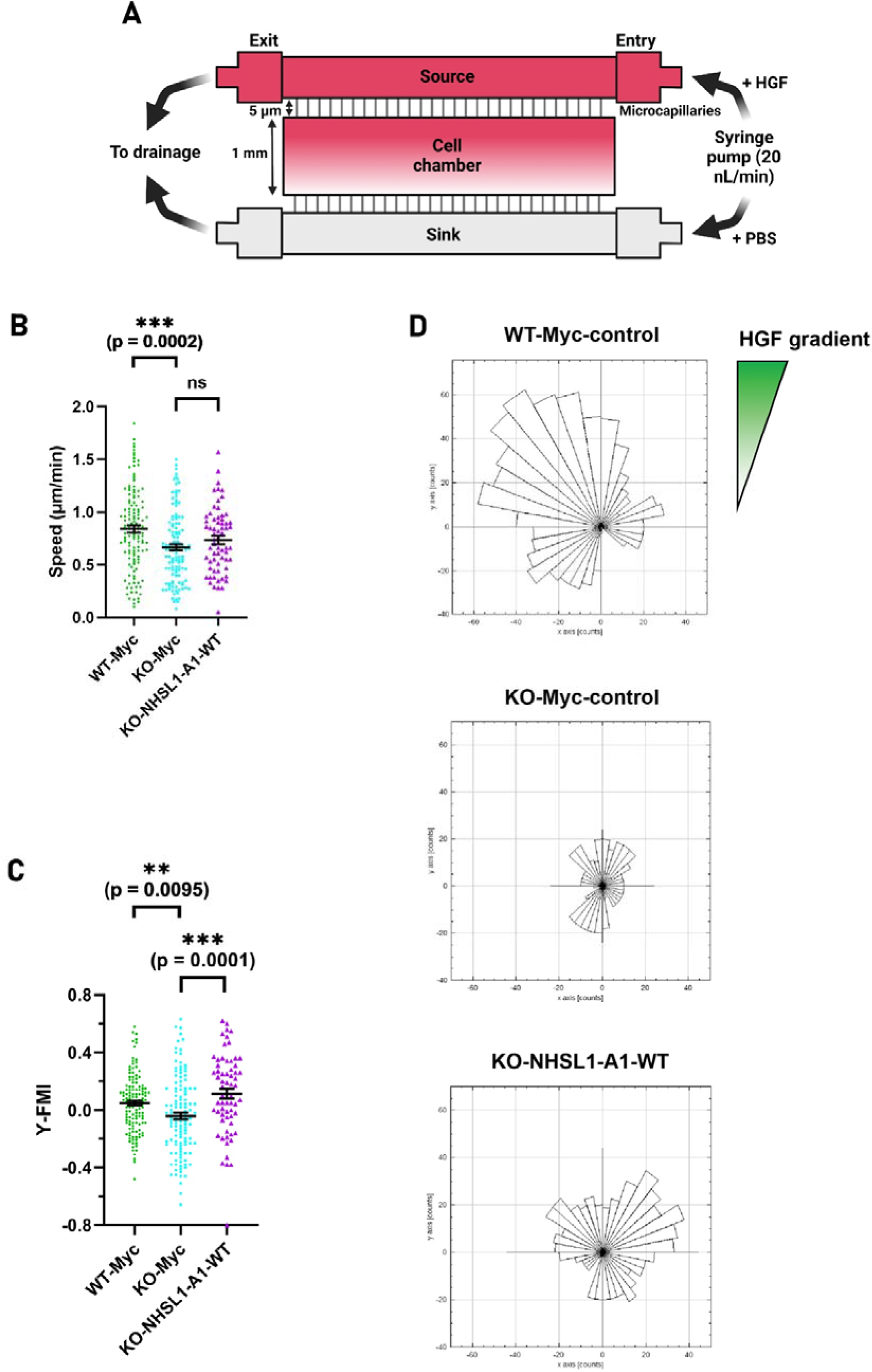
The NHSL1-A complex mediates chemotaxis. **(A)** Schematic diagram of the microfluidics chemotaxis chamber: Cells are loaded into the cell chamber through an inlet on the right side of the chamber (not depicted). Media without serum containing the chemoattractant HGF is loaded into the source inlet, while media without serum is loaded into the sink inlet. A syringe pump is then connected to both source and sink inlets to introduce a steady flow of media from source to sink in the cell chamber. **(B)** Cell migration speed and **(C)** <u>F</u>orward <u>M</u>igration Index (y-FMI) of wild-type B16F1 cells expressing the Myc-tag alone, or NHSL1 knockout B16F1 cells expressing either the Myc-tag alone or wild-type full-length NHSL1-A1-WT migrating on laminin while being subjected to a gradient of hepatocyte growth factor (HGF) in a chemotaxis chamber. **(B)** Kruskal-Wallis test: knockout vs WT control: ***P = 0.0002. **(C)** Ordinary one-way ANOVA test: knockout vs WT control: **P=0.0095; knockout vs Rescue Myc NHSL1-A-WT: ***P=0.0001. **(D**) Rose plots showing the overall direction of migration of wild-type B16F1 cells expressing the Myc-tag alone as control, or knockout B16F1 cells expressing either the Myc-tag alone as control or wild-type full-length NHSL1-A1 in a chemotaxis assay. The hepatocyte growth factor (HGF) gradient is in the direction of y-axis. Each point of origin has been reset to (x,y) = 0 on the plot, and each bar represents both the direction in which a group of cells were migrating and the number of cells (cell counts) in that group according to the respective bar length. Each bar has a range angle interval of 10° (total of 36 bars in each plot). Results are obtained from four independent biological experiments for WT control and knockout control and three independent biological experiments for Rescue Myc-NHSL1-A-WT, and n = 130 (WT control Myc only), 125 (knockout Myc only) and 65 (Rescue Myc-NHSL1-A-WT).

### The NHSL1-A complex increases lamellipodial protrusion speed via the WCA domain and reduces stability of lamellipodial protrusions

To explore the mechanisms how the NHSL1-A complex promotes cell migration, we investigated its effect on lamellipodial protrusion or retraction speeds and lamellipodial stability. To do this, we co-transfected LifeAct-mScarlet together with either EGFP as control or the EGFP-tagged NHSL1-A1 constructs and generated high magnification movies of the lamellipodium and quantified protrusion and retraction vectors at each pixel along the leading edge (Fig. 6A). This analysis revealed that the speed of lamellipodial protrusion changed from 4.7 ± 0.3 µm/min to 5.3 ± 0.4 µm/min in the NHSL1 knockout cells re-expressing wild-type NHSL1-A1, but this was not significant (Fig. 6B). Similarly, the lamellipodial protrusion speed changed from 4.7 ± 0.3 µm/min to 5.0 ± 0.3 µm/min in the NHSL1-A1-SW-Mut re-expression to a slightly lesser extent compared to NHSL1-A1-WT. However, the speed increased further to 5.9 ± 0.4 µm/min upon re-expression of the NHSL1-A1-WCA-DA and this change was significant (Fig. 6B).

**Figure 6:**
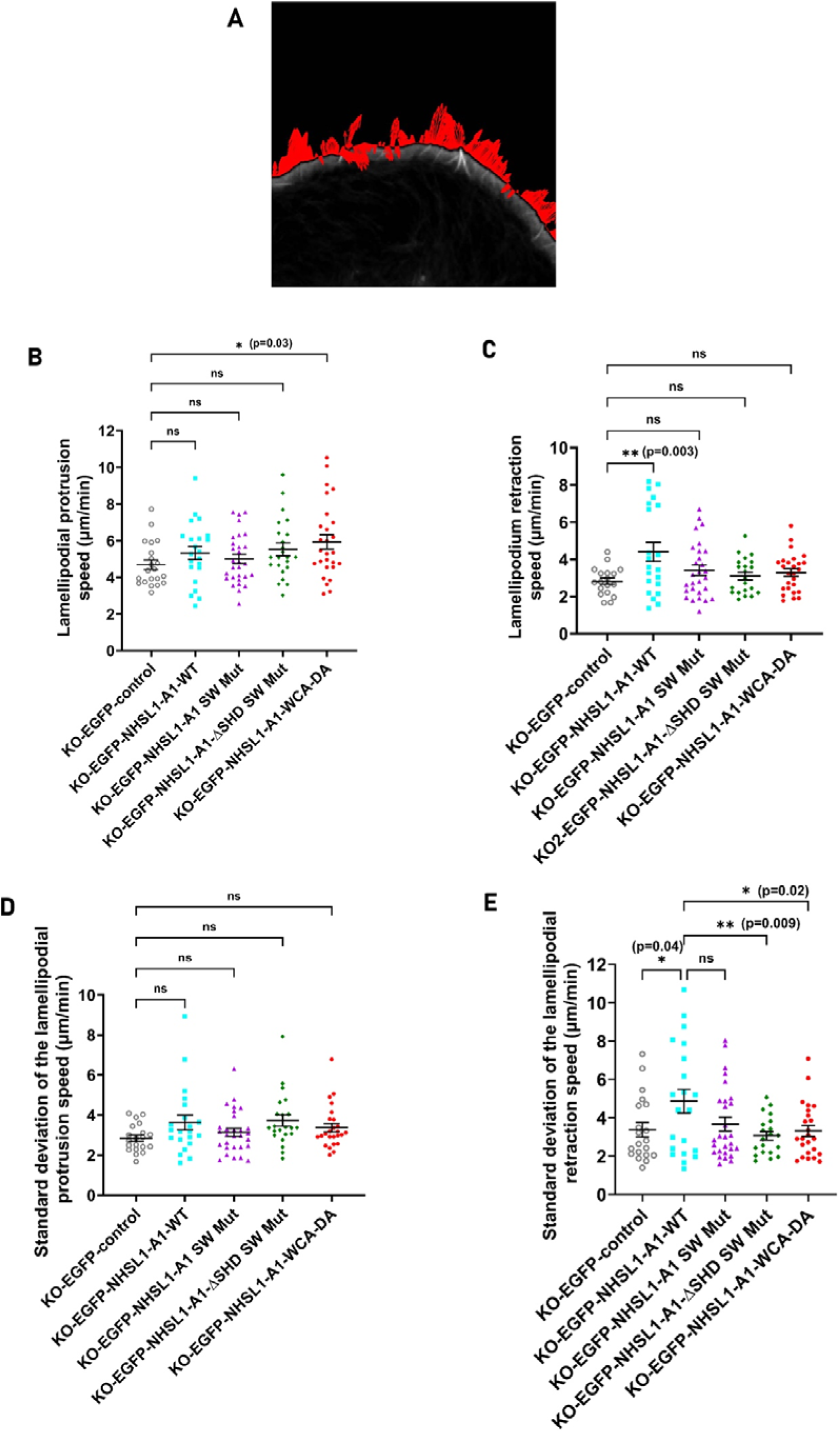
The NHSL1-A complex increases lamellipodial protrusion speed via the WCA domain and reduces stability of lamellipodial protrusions. **(A)** Example still-image from live-cell imaging of knockout B16F1 expressing EGFP only as control migrating on laminin. Cells were imaged at 160x magnification and live-cell movies were then used to quantify the lamellipodial dynamics by segmentation and calculation of protrusion and retraction vectors (see methods). **(B,C)** Quantification of lamellipodial protrusion (B) and retraction (C) speed and **(D,E)** their respective standard deviation from live-cell imaging of NHSL1 knockout B16F1 cells expressing EGFP only as control or EGFP tagged NHSL1-A constructs migrating on laminin. Results are mean ± SEM (error bars). The standard deviation of the protrusion or retraction speeds in **(D,E)** shows the fluctuations in the speed of the lamellipodial edges and hence serves as a proxy for the stability of lamellipodial protrusions. **(B)** One-way ANOVA test: NHSL1 knockout versus NHSL1 Rescue EGFP-NHSL1-A-WT: ns = 0.4827; NHSL1 knockout versus NHSL1 Rescue EGFP-NHSL1-A-SW-Mut: ns = 0.8919; NHSL1 knockout versus NHSL1 Rescue EGFP-NHSL1-A-ΔSHD-SW-Mut: ns = 0.2451; NHSL1 knockout versus NHSL1 Rescue EGFP-NHSL-1-A-WCA-DA: *P=0.03. n=21 (NHSL1 knockout EGFP only), 22 (NHSL1 knockout Rescue EGFP-NHSL1-A-WT), 29 (NHSL1 knockout Rescue EGFP-NHSL1-A-SW-Mut), 23 (NHSL1 knockout Rescue EGFP-NHSL1-A-ΔSHD-SW-Mut) and 27 (NHSL1 knockout Rescue EGFP-NHSL1-A-WCA-DA). **(C)** One-way ANOVA test: NHSL1 knockout versus NHSL1 knockout Rescue EGFP-NHSL1-A-WT: **P=0.003; NHSL1 knockout versus NHSL1 knockout Rescue EGFP-NHSL1-A-SW-Mut: ns=0.4493; NHSL1 knockout versus NHSL1 knockout Rescue EGFP-NHSL1-A-ΔSHD-SW-Mut: ns = 0.9215; NHSL1 knockout versus NHSL1 knockout Rescue EGFP-NHSL-1-A-WCA-DA: ns = 0.6599. n=18 (NHSL1 knockout EGFP only), 21 (NHSL1 knockout Rescue EGFP-NHSL1-A-WT), 28 (NHSL1 knockout Rescue EGFP-NHSL1-A-SW-Mut), 21 (NHSL1 knockout Rescue EGFP-NHSL1-A-ΔSHD-SW-Mut) and 26 (NHSL1 knockout EGFP-NHSL1-A-WCA-DA). **(D)** One-way ANOVA test: NHSL1 knockout Rescue EGFP-NHSL1-A-WT vs NHSL1 knockout: ns = 0.1223; NHSL1 knockout Rescue EGFP-NHSL1-A-WT vs NHSL1 knockout Rescue EGFP-NHSL1-A-SW-Mut: ns = 0.4167; NHSL1 knockout Rescue EGFP-NHSL1-A-WT vs NHSL1 knockout Rescue EGFP-NHSL1-A-ΔSHD-SW-Mut: ns = 0.9964; NHSL1 knockout Rescue EGFP-NHSL1-A-WT vs NHSL1 knockout Rescue EGFP-NHSL-1-A-WCA-DA: ns = 0.8728. ns. = no significant difference. n = 20 (NHSL1 knockout EGFP only), 21 (NHSL1 knockout Rescue EGFP-NHSL1-A-WT), 29 (NHSL1 knockout Rescue EGFP-NHSL1-A-SW-Mut), 22 (NHSL1 knockout Rescue EGFP-NHSL1-A-ΔSHD-SW-Mut) and 25 (NHSL1 knockout Rescue EGFP-NHSL1-A-WCA-DA). **(E)** One-way ANOVA test: NHSL1 knockout Rescue EGFP-NHSL1-A-WT vs NHSL1 knockout: *P=0.0404; NHSL1 knockout Rescue EGFP-NHSL1-A-WT vs NHSL1 knockout Rescue EGFP-NHSL1-A-SW-Mut: ns = 0.0933; NHSL1 knockout Rescue EGFP-NHSL1-A1-WT vs NHSL1 knockout Rescue EGFP-NHSL1-A-ΔSHD-SW-Mut: **P=0.0091; NHSL1 knockout Rescue EGFP-NHSL1-A-WT vs NHSL1 knockout Rescue EGFP-NHSL-1-A-WCA-DA: *P=0.0233. ns=no significant difference. n=20 (NHSL1 knockout EGFP only), 21 (NHSL1 knockout Rescue EGFP-NHSL1-A-WT), 28 (NHSL1 knockout Rescue EGFP-NHSL1-A-SW-Mut), 20 (NHSL1 knockout Rescue EGFP-NHSL1-A-ΔSHD-SW-Mut) and 24 (NHSL1 knockout Rescue EGFP-NHSL1-A-WCA-DA). Results are from four independent biological experiments.

The lamellipodial retraction speed increased significantly from 2.8 ± 0.2 µm/min in the NHSL1 knockout cells to 4.4 ± 0.5 µm/min in the knockout cells re-expressing wild-type NHSL1-A1. Retraction speed did not change in knockout cells re-expressing either NHSL1-A1 mutant variants (Fig. 6D) suggesting that the NHSL1-A complex mediates retraction speed in part via the Scar/WAVE complex. Re-expression of the NHSL1-A1-WCA-DA construct did not change the mean retraction speed (Fig. 6D). This is in contrast what we observed for this phospho-mimetic construct in protrusion speed, implying that phosphorylation-dephosphorylation cycles of the NHSL1 WCA domain may be required for its role in lamellipodial dynamics. Together, this indicates that the NHSL1-A complex may promote lamellipodial dynamics via its WCA domain and via the Scar/WAVE complex.

We then analysed the standard deviation (SD) of the mean lamellipodial protrusion or retraction speed as a readout for the lamellipodial stability over time. The SD of the lamellipodial protrusion did not change significantly between the different conditions (Fig. 6C). However, the re-expression of NHSL1-A1-WT significantly increased the SD of the lamellipodial retraction speed, but this was not the case when the NHSL1-A1-WCA-DA was re-expressed. The re-expression of the NHSL1-A1-SW-Mut may only partially increase the SD of the lamellipodial retraction speed from 3.4 ± 0.4 µm/min to 3.7 ± 0.4 µm/min (Fig. 6D). This suggests that the NHSL1-A complex reduces the stability of lamellipodial protrusions via its WCA and maybe in part via its interaction with the Scar/WAVE complex. Taken together, the NHSL1-A complex may cause lamellipodia to be faster and less stable.

### The NHSL1-A complex may activate the Arp2/3 complex in the lamellipodium

To investigate the underlying causes for NHSL1-A complex function in lamellipodia dynamics and cell migration, we quantified the Arp3 intensity in the lamellipodium (Fig. 7A,B) as a readout for the Arp2/3 complex activity as this complex is activated at the very edge of lamellipodia and then incorporated into the growing, rearward flowing F-actin branched network which it nucleates^1^. We quantified Arp3 intensity in the lamellipodium in NHSL1 knockout cells re-expressing EGFP-tagged NHSL1-A1 constructs. We also quantified the cellular EGFP levels and used it to normalise the Arp3 intensity to the expression level of the rescue constructs on a cell-by-cell basis. We observed that lamellipodial Arp2/3 levels in NHSL1 knockout cells upon re-expression of wild-type NHSL1-A1 increased by 20% but upon re-expression of NHSL1-A1-SW-Mut decreased by 12% but these changes were not significant. In contrast, the re-expression of NHSL1-A1-ΔSHD-SW-Mut caused a significant reduction of 40% whereas re-expression of NHSL1-A1-WCA-DA did not change Arp2/3 intensity in the lamellipodium (Fig. 7B). This suggest that the NHSL1-A complex may promote Arp2/3 activation in the lamellipodium via the Scar/WAVE complex.

**Figure 7:**
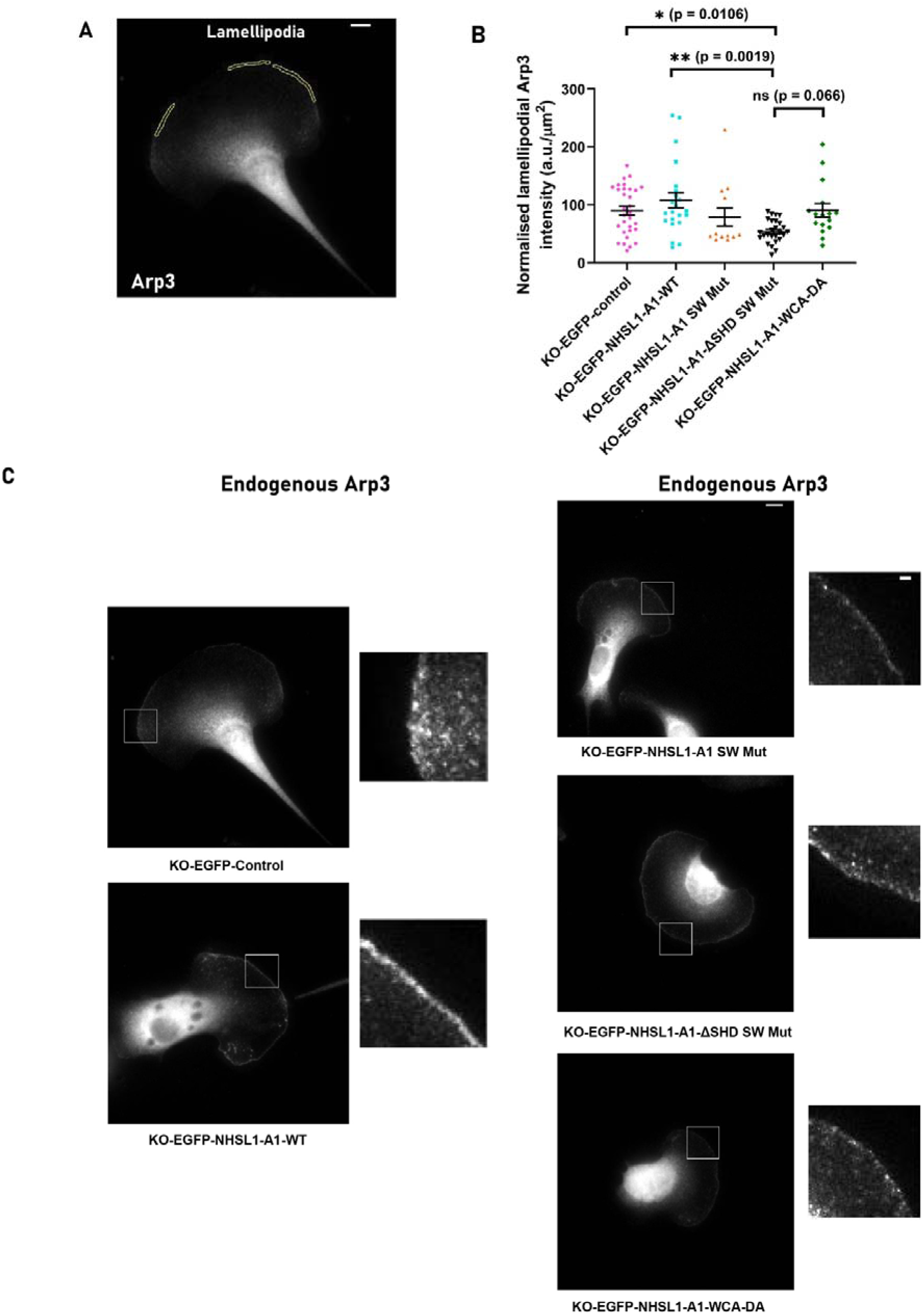
The NHSL1-A complex may increase Arp2/3 activity in the lamellipodium. **(A)** Example of a B16F1 cell with an outline of the lamellipodium for measurement of Arp3 intensity. **(B)** Quantification of normalised Arp3 intensity in the lamellipodium; Kruskal-Wallis test: NHSL1 knockout versus NHSL1 knockout Rescue EGFP-NHSL1-A1-WT: ns > 0.9999; NHSL1 knockout versus NHSL1 knockout Rescue EGFP-NHSL1-A1-SW-Mut: ns > 0.9999; NHSL1 knockout versus NHSL1 knockout Rescue EGFP-NHSL1-A1-ΔSHD-SW-Mut: *P = 0.0106; NHSL1 knockout versus NHSL1 knockout Rescue EGFP-NHSL1-A1-WCA-DA: ns > 0.9999; NHSL1 knockout Rescue EGFP-NHSL1-A1-WT versus NHSL1 knockout Rescue EGFP-NHSL1-A1-SW-Mut: ns > 0.9999; NHSL1 knockout Rescue EGFP-NHSL1-A1-WT versus NHSL1 knockout Rescue EGFP-NHSL1-A1-ΔSHD-SW-Mut: **P = 0.0019; NHSL1 knockout Rescue EGFP-NHSL1-A1-WT versus NHSL1 knockout Rescue EGFP-NHSL1-A1-WCA-DA: ns > 0.9999; NHSL1 knockout Rescue EGFP-NHSL1-A1-SW-Mut versus NHSL1 knockout Rescue EGFP-NHSL1-A1-ΔSHD-SW-Mut: ns > 0.9999; NHSL1 knockout Rescue EGFP-NHSL1-A1-SW-Mut versus NHSL1 knockout Rescue EGFP-NHSL1-A1-WCA-DA: ns > 0.9999; NHSL1 knockout Rescue EGFP-NHSL1-A1-ΔSHD-SW-Mut versus NHSL1 knockout Rescue EGFP-NHSL1-A1-WCA-DA: ns = 0.0666. Results are obtained from four independent biological experiments, and n = 32 (NHSL1 knockout), 22 (NHSL1 knockout Rescue EGFP-NHSL1-A1-WT), 15 (NHSL1 knockout Rescue EGFP-NHSL1-A1-SW-Mut), 26 (NHSL1 knockout Rescue EGFP-NHSL1-A1-ΔSHD-SW-Mut) and 16 (NHSL1 knockout Rescue EGFP-NHSL1-A-WCA-DA). **(C)** Example images of B16F1 NHSL1 knockout cells re-expressing various EGFP-NHSL1-A1 constructs or EGFP as control and stained for endogenous Arp3. Representative images from four independent experiments. Scale bar represents 10 µm. Inset is a magnified image of the white box and scale bar represents 2 µm.

This increased lamellipodial Arp2/3 mediated F-actin nucleation may increase lamellipodia protrusion speed and consequently, increase cell migration speed.

## Discussion

We have shown here that an NHSL1-A complex exists whose formation is mediated by the SHD domain of the NHSL1-A isoforms. Expression of only the NHSL1-A1 isoform in pan-isoform NHSL1 knockout cells induced dynamic lamellipodia and aster like vesicular compartments from which vesicles appear to bud off, both locations at which NHSL1-A1 resides. We identified a functional WCA domain in all NHSL1 isoforms whose interaction with the Arp2/3 complex is promoted by GSK3 phosphorylation. Re-expression of NHSL1-A1 with dominant active mutations in its WCA domain increased Arp2/3 interaction, lamellipodial protrusion but not retraction speed nor decreased lamellipodial stability suggesting that the WCA domain of NHSL1 promotes productive lamellipodial protrusion. In agreement, re-expression of NHSL1-A1 with dominant active mutations in its WCA domain increased migration speed but not the NHSL1 SHD on its own nor NHSL1-A1 mutated in its Scar/WAVE complex binding sites suggesting that the NHSL1-A complex functions to promote migration efficiency via both its WCA domain and the Scar/WAVE complex.

Our study revealed that the SHD in NHSL1-A proteins is fully functional and mediates the formation of a complex which resembles the Scar/WAVE complex but in which Scar/WAVE1-3 proteins are replaced by NHSL1-A. In agreement, the SHD of the related NHS protein was found to bind to Scar/WAVE complex components^21^. While our study was underway Wang et al reported that the PPP2R1A subunit of the PP2A phosphatase specifically interacted with the NHSL1-A1 isoform and components of the Scar/WAVE complex but not Scar/WAVE proteins ^24^. This was surprising as NHSL1-A1 has two Abi binding sites mediating interaction with the entire Scar/WAVE complex suggesting that the NHSL1-Scar/WAVE complex interaction is tightly regulated. In addition, they found that the SHD of NHSL1-A interacted with *in vitro* translated Scar/WAVE components^24^ which supports our findings. Interestingly, knockout of the Scar/WAVE complex components Nap1 or CYFIP1/2 have been widely used to study the function of the Scar/WAVE complex^15,41–46^. In light of our findings that these also constitute components of an NHSL1-A complex it is possible that the reported defects may be caused by the loss of the NHSL1-A complex and/or combined loss of both complexes.

We found that the NHSL1-A complex localises to the very edge of lamellipodia and to vesicles similarly to what we have previously shown for the NHSL1-F1 isoform which does not harbour an SHD^25^. NHSL1-A1’s recruitment to lamellipodia depended on the SHD but not on the interaction with the Scar/WAVE complex suggesting both complexes are independently recruited. However, its recruitment to vesicles does not depend on the SHD which is in agreement with our findings that fragment 1 and 3 of NHSL1-F1 mediate vesicle localisation^25^. The NHSL1-F1 isoform does not contain an SHD and yet localises to the leading edge presumably being recruited there by its two Rac binding sites^25^. The NHSL1-A1 Scar/WAVE binding mutant displayed much weaker leading edge and vesicles localisation suggesting that its localisation at leading edge and vesicles is stabilised by its interaction with the Scar/WAVE complex but is not mediated by it. Interestingly, we observed that this construct also localised to the tips of microspikes and filopodia suggesting that this interaction restricts it to lamellipodia and that the NHSL1-A complex may have an additional, regulated unknown function at this location.

Because the Scar/WAVE proteins mediate the function of the Scar/WAVE complex via its C-terminal WCA domain by recruiting and activating the Arp2/3 complex, we speculated that NHSL1 may have a similar function in the NHSL1-A complex. Indeed, we discovered a functional WCA domain in all NHSL1 isoforms which mediates interaction with the Arp2/3 complex. The interaction between the NHSL1 WCA and the Arp2/3 complex is promoted by GSK3 phosphorylation. It was previously unknown that GSK3 phosphorylation increases WCA-Arp2/3 interaction, but we found that GSK3 phosphorylation may also increase the Scar/WAVE-WCA Arp2/3 interaction (Fig. 2B-D). Fittingly we recently showed that GSK3 activity is required for lamellipodia formation^47^.

It has been shown that two nucleation promotion factors with WCA domains have to be present for full activation of the Arp2/3 complex^48,49^. This raises the exciting possibility that the Scar/WAVE-WCA and the NHSL1-WCA cooperate to activate the Arp2/3 complex because the Scar/WAVE and NHSL1-A complexes interact at the leading edge.

We showed that re-expression of NHSL1-A1 with phospho-mimetic, dominant-active mutations in the WCA domain significantly increased lamellipodial protrusion speed but not retraction speed and also did not decrease lamellipodial stability suggesting that the WCA domain of NHSL1 promotes productive lamellipodial protrusion and thus cell migration efficiency. This also suggests that the NHSL1 WCA is partially functional but needs to be dephosphorylated like the Scar/WAVE2 WCA to activate the Arp2/3 complex.

Re-expression of wild-type NHSL1-A1 also appeared to increase lamellipodial protrusion speed albeit to a lesser extent. In agreement, knockdown of NHSL1b in zebrafish with a morpholino targeting the SHD containing exon 1 caused reduction in lamellipodial protrusion speed and persistence^50^. In contrast, we previously showed that knockout of all NHSL1 isoforms, increased stability of lamellipodial protrusions without affecting protrusion speed while rescue with the NHSL1-F1 isoform reduced lamellipodial protrusion stability^25^ suggesting that the NHSL1-F1 isoform has a different function compared to the NHSL1-A complex.

Similarly, we showed that knockout of all NHSL1 isoforms increased cell migration speed and persistence in 2D random cell migration assays^25^ and this has been confirmed with a knockdown in another cell line^24^. When we re-expressed wild-type NHSL1-A1 in an NHSL1 knockout cell line this increased migration speed via the Scar/WAVE complex. Also, re-expression of NHSL1-A1 with a dominant active WCA mutation increased migration speed like wild-type but not re-expression of the NHSL1-A SHD on its own suggesting that the complex functions to promote migration via both the Scar/WAVE complex and the WCA domain. Recently we reported that NHSL1 promotes Fast Endophilin Mediated Endocytosis (FEME) in cells and its interactions with Endophilin-A2 and Ena/VASP proteins are required for this function^51^. Thus, NHSL1 function in endocytosis may contribute to its effect on migration efficiency since re-expression of NHSL1-A1-ΔSHD-SW-Mut increased migration speed like wild-type NHSL1-A.

We found that the NHSL1-A complex is required for chemotaxis and NHSL1 knockout cells displayed a reduced migration speed during chemotaxis. In contrast, knockdown of all NHSL1 isoforms increased the ability of cells to migrate up a gradient of fibronectin showing that NHSL1 functions differently in chemotaxis versus haptotaxis^24^. Escot et al also observed a reduction in directional migration upon knockdown of the SHD containing isoform of NHSL1b in zebrafish with a reduced cell migration speed and persistence^50^ suggesting that this directed migration may be due to chemotaxis. Zebrafish NHSL1b was also required for a different type of directed migration, the facial branchiomotor (FBM) neuronal migration in a planar-polarised environment^52^. As the Arp2/3 complex is not required for chemotaxis^40,53^, NHSL1 may control chemotaxis in an Arp2/3 complex independent manner either via an Arp2/3 complex independent Scar/WAVE complex function or via its role in endocytosis.

Taken together, this suggests that the regulation of actin nucleation by the Arp2/3 complex is much more complex than anticipated.

## Methods

### Molecular Biology and Plasmids

Full-length human NHSL1-A1 wild-type and NHSL1-A1-SW-Mut cDNA was generated via HiFi cloning (NEB Hifi, New England Biolabs, Inc.) by adding the Scar Homology Domain (SHD) sequence from exons 1 and 2 (Geneblock: CTTTAAAGGAACCAATTCAGTCGACTGGATCCGGACCGAATTCgccaccATGCCCTTCCACCAgaGgAcCcTgGAGCCCGCgCGGCTGCGCCGGCCCGAGGCGGCgggGGCcgGGGCTGcGGgCgCGCCaCTCTTCCGCTCGCTGGAGCAGGTCAGCTCGCAtgCCCTGGgCTGCCTGCTgGCgCAGCTGGCCGACCTGTCGCGCTGCGCcGGGGACATcTTCGGcGAGCTCGAGGGCCAGGCGGCcGCGCTGGGcCACCGCACCGCCGcGcTGCACCGcCGCCTcGACGCCCTGCAaGCCGCCGCtGCGCGcCTGGACCACCGcCGAGTGAAAATCCCGGTTTCCAACCTAGACTGCCCCGTGGCACCAGCAGGAGAATGTGTTCCTTCCCACCACAAGACCCCCCTGTGTTGAGGACCTGCACCGCCAAGCCAAGCTCAACCTCAAATCAGTACTGAGGGAATGCGATAAGTTGCGGCATGATGGCTACCGCAGTTCTCAGTACTACTCTCAGGGACCCACCTTTGCGGCCAACGCCAGCCCATTCTGTGATGATTACCAAGATGAAGATGAAGAAACAGATCAAAAGTGTTCTCTGTCTTCATCAGAAGAAGAAAGATTTATTTCCATCAGGAGACCTAAAACACCAGCCTCAAGTGACTTCTCCGACCTTAATACTCAGACGAACTGGACCAAGTCGCTTCCACTG) of NHSL1-A to the 5’ end of NHSL1-F1. To generate the SHD of NHSL1-A1 alone, it was amplified from NHSL1-A1 using 5’-primer: cgGGATCCGCCACCATGCCATTTCACCAG and 3’-primer: cgGAATTCgcCTCGCGGAGAACGGATTTAAGG and cloned into the pENTR3C plasmid (Invitrogen) using restriction enzyme-based cloning. NHSL1-A1 in which the SHD was deleted (exon 1 and first few bases of exon 2 removed) was generated by amplifying it from full-length NHSL1-F1-SW-Mut using 5’-primer: cggGGATCCgccaccatgGTTGAGGACCTGCACCGCC and 3’-primer: cggTCTAGAcgACTCTCCTCGCTCAGAGAACC. Full-length NHSL1-A1 with a dominant active WCA domain was constructed by HiFi cloning a synthetic double-stranded DNA (Geneblock, IDT) incorporating phosphomimentic mutations within the WCA (ATCAATGTTTTTGTTGGAAGAGCTCAGAAAAACCAAGGGGACCGGTCCAATTACCAGGATAAATCCCTATCAAGAAACATCTCTTTGAAGAAAGCAGCGAAGGCTGCCCTGCCAGCCTCGCGGACAGACTCCCTCCGCAGGATTCCCAAGAAGAGCAGCCAGTGCAACGGGCAGGTGCTCAACGAGAGCCTGATCGCCACACTCCAGCACTCGCTGCAGCTGAGCCTCCCAGGCAAAAGTGGCAGCGAACCCTCCCAGGAACCCTGCGAAGACTTGGAAGAGCCCTGGCTGCCCCGCTCCCGGAGCCAGAGCACAGTTAGTGCTGGCAGCAGCATGACTTCCGCCACCACCCCCAATGTCTACTCCCTGTGCGGGGCCACGCCATCGCAGAGTGACACAAGCAGCGTCAAGTCAGAGTACACGGACCCCTGGGGTTATTACATTGACTACACGGGCATGCAGGAAGATCCGGGGAACCCGGCAGGGGGCTGTTCAACCAGCAGTGGGGTGCCCACTGGGAACGGGCCAGTCCGCCATGTCCAAGAAGGGTCCAGAGCCACAATGCCCCAAGTGCCCGGTGGTTCAGTCAAACCAAAGATCATGTCACCAGAGAAGTCACACAGAGTCATTTCTCCATCCAGTGGGTATTCCAGCCAGTCGAATGCACCCACAGCACTCACCCCTGTGCCTG TGTTTT) using full-length NHSL1-A1 in pENTR3C as the backbone. GST-tagged WCA wild-type and dominant active were generated by amplifying the WCA region of NHSL1-A with primers (for wild type, 5’-primer: cggGgatccgccaccatgGTTGGAAGAGCTCAGAAAAACC and 3’-primer: cggTCTAGAcgCCAtcaACCTCCTCCTCCTCCCAGCCAGGGCTCTTCCAAG; for dominant active, 5’-primer: cggGgatccgccaccatgGTTGGAAGAGCTCAGAAAAACC and 3’-primer: cggTCTAGAcgCCAtcaACCTCCTCCTCCTCCCCAGGGCTCTTCCAAGTCTTC). GST-tagged WCA of WAVE2 was constructed by amplifying the region of Scar/WAVE2 with 5’-primer: cgcGGATCCgccaccatgagcgatgcccgtagcgacc and 3’-primer: cggTCTAGAccgCCTCCTCCTCCatcggaccagtcgtcctcatc. The tagged Scar/WAVE complex components were described in^25^. GSK3 cDNA was amplified and cloned into pENTR3c by restriction cloning. The following Gateway (Invitrogen)compatible mammalian expression plasmids were cloned by traditional restriction-based cloning or Gibson assembly (NEB Hifi, New England Biolabs, Inc.): pCAG-EGFP-DEST, pCAG-DEST-EGFP, pDEST15, pCAG-EGFP-DEST-IRES-Puro, pRK5Myc-DEST, pCAG-Myc-DEST, pCAG-Myc-DEST-IRES-BLAST, and pStrep-tagII-DEST. cDNAs in pENTR3C were transferred into these tagged mammalian expression vectors using the Gateway recombination method (Invitrogen).

### Antibodies

The NHSL1 rabbit polyclonal antibody recognising all NHSL1 isoforms (Eurogentec, #4457) has been described previously^25^. The NHSL1-A SHD specific antibody was generated against a mixture of two peptides (Eurogentec): Human (aa6-20: C+RTLEPARLRRPEAAG) and Mouse (aa6-20: C+RSVEPARLRRPDEAA). Commercial primary antibodies: EGFP (Roche; cat 11814460001: (mouse mAb IgG1k; mixture of clones 7.1 and 13.1)), Myc (9E10, Sigma), Abi1 (MBL, clone 1B9, D147-3), Scar/WAVE1 mAb (BD 612276), Arp3 (Cell Signaling 4738 or Abcam ab151289), Scar/WAVE2 rabbit mAb (D2C8, CST 3659), HSC70 (Santa Cruz sc7298). Secondary antibodies: HRP-anti-rabbit (7074S), HRP-anti-mouse (7076S) (Cell Signaling Technology). AlexaFluor-Plus488-goat anti rabbit IgG (Invitrogen A32731), AlexaFluor488 goat anti mouse IgG (Invitrogen A11029), AlexaFluor568 goat anti mouse IgG (Invitrogen A11004), AlexaFluor568 goat anti rabbit IgG (Invitrogen A11036).

### Glutathione S-transferase (GST) and Strep-tag-II purification

BL21(DE3)-RP bacteria (Stratagene) were transformed with a DNA construct of interest by heat-shock treatment and colonies were grown on agar plates containing specific antibiotics. Single colonies were grown in LB medium, 200 ug/ml ampicillin, and 2% D-glucose until OD_600_ = 0.5. Protein expression was induced by adding 0.1mM Isopropyl β-d-1 thiogalactopyranoside (IPTG) into the bacterial culture and incubated over night at 18C. The culture was harvested at 8000 x g for 20 minutes at 4°C, resuspended in 10 ml ice-cold PBS with protease inhibitors (Roche tablets Mini without EDTA), and pulse sonicated on ice for 3 minutes at 15 Watts before it was centrifuged again at 18,000 x g for 20 minutes at 4°C. The supernatant was incubated with Glutathione Sepharose™ 4 Fast Flow Media (GE Healthcare) or Streptactin-XT (IBA Life Sciences, Germany) for 2 hours at 4°C, washed with cold PBS. Purity was assessed by a Coomassie staining of SDS-PAGE gels.

### Cell cultures and transient transfection

HEK293FT cells (Thermo Fisher Scientific R70007), B16-F1 mouse melanoma cells (ATCC CRL-6323), and the B16-F1 NHSL1 knockout cell line^25^ were cultured in Dulbecco’s modified Eagle’s medium containing 1% penicillin/streptomycin, 2 mM L-glutamine and 10% fetal bovine serum (Gibco). The B16-F1 NHSL1 knockout cell line (clone 2) was generated and characterised as described previously by knocking in a stop codon into exon 2 common to all NHSL1 isoforms^25^. HEK293FT cells were transiently transfected using PEI (Sigma 913375 or 408727: 2ug DNA in 100ul OptiMEM to 4ul 2mg/ml PEI in 100ul OptiMEM, mixed, and incubated for 20 min at room temperature). B16-F1 cells were transfected with XtremeGENE™ 360 transfection reagent (Roche) according to the manufacturer’s instructions and replaced with normal growth media after 4–6 hours. For random migration and chemotaxis assays, B16-F1 cells transfected with a bicistronic expression plasmid were selected with blasticidin for 48 hours before plating to ensure that all cells analysed were expressing the specific protein of interest. All cells (B16-F1, HEK 293FT) were maintained at 37°C in 10% CO2.

### Immunoprecipitation, pulldown assay and Western blotting

Cells were lysed with glutathione S-transferase (GST) buffer (50 mM Tris-HCL, pH 7.4, 200 mM NaCl, 1% NP-40, 2 mM MgCl_2_, 10% glycerol, 10 mM NaF, 1 mM Na_3_VO_4_, and complete mini tablets without EDTA (Roche)). Lysates were incubated on ice for 15 minutes and centrifuged at 17,000 × g at 4°C for 10 minutes. Protein concentration was then determined using Precision Red Advanced Protein Assay (Cytoskeleton Inc.).

For co-pulldown assays, the lysates were incubated with either GFP-selector beads (Nano Tag Biotechnology) or Myc-trap beads (Chromotek). The beads were blocked with 1% BSA in GST buffer before incubating with lysates for 1–2 hours. Following this, the beads were washed in GST buffer three times before boiling in 1x gel sample buffer (5x sample buffer: 200 mM Tris-HCl pH6.8, 5% w/v SDS, 25% v/v glycerol, 233 mM DTT, 0.075% (w/v) bromophenol blue) for 5 minutes. The beads were separated on SDS-PAGE gels (10x buffer: 250mM Tris-Base,1.92M Glycine,1%(w/v) SDS) and transferred onto Immobilon-P membranes (EMD Millipore). Western blotting was performed by transferring at 100 V, 350 mA, 50 W for 1.5 hours before blocking in 5% (w/v) BSA or 5% (w/v) milk in 1X TBS-T (diluted from 10x TBS-T: 200 mM Tris (pH7.6), 1.54 M NaCl, 1% (v/v) Tween-20) overnight. The blots were then incubated with the indicated primary antibodies for 1 hour followed by HRP conjugated secondary antibodies (Cell Signalling Technology) for 1hour at room temperature with three washes of TBST, TBST+0.5 M NaCl and TBST+0.5% (v/v) Triton X-100 (TX-100). Blots were developed with Clarity Western ECL Substrate (Bio-Rad Laboratories) using a Bio-Rad Imager and ImageLab software. Blots were quantified using the ImageLab software by measuring pixel intensity of the desired protein bands and normalising them to their corresponding cell lysate pixel intensity.

### Live-cell imaging and quantification of lamellipodial dynamics

B16-F1 cells were plated in DMEM, high glucose, 2 mM glutamine, no phenol red (Gibco) onto a 35 mm glass-bottom µ-Dish (Ibidi) coated with 25 µg/mL laminin (L2020, Sigma) or 10 µg/ml fibronectin (Sigma; F1141) 3–4 hours before imaging. An IX81 microscope (Olympus) with a Solent Scientific incubation chamber, filter wheels (Sutter), an ASI-XY stage, Cascade-II 512B camera (Photometrics), and 4× UPlanFL 0.13, 10× UPlanFL 0.30, 60x Plan-Apochromat NA 1.45, or 100× UPlan-Apochromat S NA 1.4 objective lenses controlled by Meta Morph software was used for live imaging. Images were taken every 10 seconds for 10 minutes at 100x or 160x magnifications using a 1.6x intermediate magnification lens (Olympus IX81), and equal exposure times were used for all cells imaged within the same condition for a given experiment. Each movie was subjected to pre-processing using FIJI (ImageJ), which included denoising, background removal and normalised contrast enhancement. Denoising was carried out using the PureDenoise plugin from the Bioimaging Group (BIG) at EPFL (available from https://bigwww.epfl.ch/algorithms/denoise/) which implements a mixed Poisson-Gaussian noise model. Automatic global noise estimation was implemented with 10 cycle spins and 3 adjacent frames used to estimate each frame. After processing, the movies were analysed using a MatLab script kindly provided by Andrew Jamieson and Gaudenz Danuser (UT Southwestern, USA) and available at: https://github.com/DanuserLab/Windowing-Protrusion. The lamellipodial protrusion speed and lamellipodial stability (standard deviation of the lamellipodial speed) of randomly migrating B16-F1 cells plated on laminin were quantified by automatically segmenting the cell outline in each frame of the movies and calculating protrusion vectors at each pixel along the cell edge. The standard deviation of the lamellipodial speed shows the fluctuation of speeds along the edge and serves as a measure for the stability of lamellipodial protrusions.

### Random migration assay and quantification

B16-F1 cells were plated onto 25ug/ml laminin-coated 4-well µ-Slide (Ibidi) in DMEM, high glucose, no glutamine, no phenol red (Gibco, Thermo Fisher) for 2–3 hours before imaging for 16 hours every 5 minutes. Cells were manually tracked by their nuclear position using the Manual Tracking plugin (FIJI), and the cell track coordinates were imported into Mathematica for analysis using the Chemotaxis Analysis Notebook v1.6γ (G. Dunn, T. Pallett, R. Marsh, King’s College London, UK).

Speed measurements from tracking data are susceptible to positional error (due to manual tracking) as well as biological noise (from cell morphology changes or nuclear repositioning). To address the former, we estimated the positional error by tracking the same cell multiple times (each time blinded to the previous track) and then calculated the time interval (as an integer multiple of the frame interval) over which this error fell to at least 10% of the average displacement measurement (this value was 10 min, or 2 frames). Biological noise can be estimated from looking at the peak of the directionality ratio, which corresponded to approximately 20 min (or 4 frames). We selected the smallest multiple for which these conditions were met (i.e. 20 min) to avoid under-sampling the instantaneous path length. Instantaneous cell speed is the displacement over the usable time interval; the Mean Track Speed (MTS) is then the average cell speed over the whole track length.

The directionality ratio over time and mean square displacement were calculated using Microsoft Excel macros provided in^54^ and were run according to the instructions provided. The directionality ratio was normalised between 0 (corresponding to the expected directionality ratio for a pure random walk over that time interval) and 1 (corresponding to a straight line). The alpha values (which characterise the motion as described in the main text) are calculated from the gradient of the log-log transform of the MSD, determined in this case by fitting a straight-line to the transformed data (from a time interval of 20 min up to 120 min) using (linear) unweighted least-squares regression. The lower interval of 20 min was chosen to match the interval used in the calculation of MTS, which avoids fitting the positional error and biological noise. The upper interval of 120 min was chosen conservatively to avoid fitting to longer time intervals for which the log-log plot deviates from a straight line.

### Microfluidics chemotaxis assay

#### Microfluidics device preparation

The pattern for the chemotaxis chamber was fabricated on silicon wafers using a photolithography process by TE connectivity (microLiquid, Spain), and the silicon wafer was exposed to silane overnight. Polydimethylsiloxane (PDMS) made from SYLGARD™ 184 Silicone Elastomer Kit (Dow Corning) was then poured on the wafer and cured overnight at 37°C. Using forceps and a razor blade, individual PDMS devices were cut from the wafer and stored in a clean dish. Holes were punched with Rapid Core 0.75 (WellTech) to create ports on the inlets and outlets of the source and sink channels and the cell chamber. The devices were washed with 70% ethanol and blow-dried, while a 35 mm µ-Dish (Ibidi) was washed with deionised water and 70% ethanol and blow-dried. Both the PDMS device and the dish were then exposed to oxygen plasma for 2 minutes. The PDMS device was then immediately placed into contact with the dish bottom to ensure an irreversible bond was formed.

#### Chemotaxis assay

B16-F1 cells in DMEM, high glucose, HEPES, 2mM Glutamine, no phenol red (Gibco; 21063029) were loaded onto the cell chamber of the PDMS device using a gel loading pipette tip. The outlet port of both source and sink channels were connected to an Eppendorf tube via a PEEK tubing (1579 Tube, Peek, NAT 1/32 x 0.0025 x 5FT, WO: 704609) (IDEX Health and Sciences). 100 µl Hamilton glass syringes (81020,1710TLL 100 µl SYR) were connected to 27 ½ gauge needles connected to PEEK tubings. The source syringe and tubing were filled with serum-free DMEM containing 100ng/mL hepatocyte growth factor (HGF) with 10µg/mL Dextran, Oregon Green™ 488 (Thermo Fisher Scientific) to visualise the chemotactic gradient. The sink syringe and tubing were filled with serum-free DMEM. The tubings were then inserted into the inlets of source and sink channels respectively, and the syringes were placed and secured on a World Precision Instruments Microfluidics Double syringe pump (NE-4002X-BS) before turning the pump on at a flow rate of 20 nL/min. A stable chemotactic gradient was then visually confirmed after 1 hour by measuring the fluorescence intensity of the Oregon Green labelled dextran along a straight line, drawn from the top to bottom of the cell chamber in the Metamorph software.

#### Directional migration acquisition and analysis

Chemotaxis assays were performed on an IX81 Olympus microscope (see above) at 10x magnification using MetaMorph imaging software. Images were taken every 10 minutes for 16 hours. Cells were manually tracked using the FIJI (ImageJ) software Manual Tracking plugin. The tracks were then analysed using the ImageJ Chemotaxis and Migration Tool (Ibidi: https://ibidi.com/chemotaxis-analysis/171-chemotaxis-and-migration-tool.html). Forward migration index in the direction of the chemotactic gradient (y-FMI) and speed were extracted with this tool alongside rose diagrams to show the direction of cell migration.

### Statistical analyses

Sample data were tested for normal distribution by D’Agostino-Pearson and Shapiro-Wilk normality tests. Based on the normality test results, the sample size, variance equality, and reasonable expectation of a normal distribution, statistical analysis was performed in Prism v7-10 (GraphPad Software) using Student’s t-test, One-way ANOVA or non-parametric Kruskal–Wallis test with appropriate corrections (e.g. Welch’s correction for unequal variance) and post-hoc tests (see figure legends in each case). P values < 0.05 were considered significant.

## Acknowledgments

We thank Andrew Jamieson and Gaudenz Danuser (UT Southwestern, USA) for providing the Windowing MATLAB script and R. Marsh, King’s College London, UK for help with Mathematica. S.J. was supported by a Malaysian Public Service Department (PSD) studentship. T.P. was supported by an Engineering and Physical Sciences Research Council (EPSRC) studentship. This work was supported by grants from the Biotechnology and Biological Science Research Council (BBSRC), UK (BB/N000226/1; BB/R015953/1) (M.K.) and from the National Institute of Health (NIH), United States (R35 GM130312/GM/NIGMS) (J.B.).

## Declaration of interests

The authors declare no competing interests.

## Supplemental movie legends

**Supplemental movie 1: EGFP-control does not localise lamellipodia**

This movie shows that EGFP expressed as control together with LifeAct-mScarlet-I in B16-F1 cells plated on laminin does not localise to the edge of protruding lamellipodia. The LifeAct-mScarlet-I serves as a marker for F-actin. Representative movie shown from three independent biological repeats. Cells were imaged every 10 seconds for the indicated times in minutes and seconds by wide-field time-lapse video microscopy using an IX 81 microscope (Olympus). Scale bar: 20 mm.

**Supplemental movie 2: EGFP-NHSL1-A1 localise to the edge of lamellipodia and vesicles**

This movie shows that EGFP-NHSL1-A1-WT expressed together with LifeAct-mScarlet-I in B16-F1 cells plated on laminin localises to the edge of protruding lamellipodia and to vesicles. The LifeAct-mScarlet-I serves as a marker for F-actin. Representative movie shown from three independent biological repeats. Cells were imaged every 10 seconds for the indicated times in minutes and seconds by wide-field time-lapse video microscopy using an IX 81 microscope (Olympus). Scale bar: 20 mm.

**Supplemental movie 3: EGFP-NHSL1-A1 in addition localise to F-actin-rich “aster” structures from which vesicles appear to bud off**

This movie shows that in around half of cells EGFP-NHSL1-A1-WT expressed together with LifeAct-mScarlet-I in B16-F1 cells plated on laminin localises to F-actin-rich “aster” structures from which vesicles appear to bud off in addition to its lamellipodia localisation. The LifeAct-mScarlet-I serves as a marker for F-actin. Representative movie shown from three independent biological repeats. Cells were imaged every 10 seconds for the indicated times in minutes and seconds by wide-field time-lapse video microscopy using an IX 81 microscope (Olympus). Scale bar: 20 mm.

**Supplemental movie 4: EGFP-NHSL1-A1-SW-Mut localise weakly to lamellipodia and some vesicles but in addition to the tips of microspikes and filopodia**

This movie shows that EGFP-NHSL1-A1-SW-Mut expressed together with LifeAct-mScarlet-I in B16-F1 cells plated on laminin localises weakly to the edge of lamellipodia and some vesicles. In addition, in contrast to NHSL1-A1-WT, it also localises to the tips of miscrospikes and filopodia. The LifeAct-mScarlet-I serves as a marker for F-actin. Representative movie shown from three independent biological repeats. Cells were imaged every 10 seconds for the indicated times in minutes and seconds by wide-field time-lapse video microscopy using an IX 81 microscope (Olympus). Scale bar: 20 mm.

**Supplemental movie 5: EGFP-NHSL1-A-ΔSHD-SW-Mut localises to the cytoplasm and weakly to some vesicles**

This movie shows that **EGFP-NHSL1-A-**Δ**SHD-SW-Mut** expressed together with LifeAct-mScarlet-I in B16-F1 cells plated on laminin localizes to the cytoplasm and weakly to some vesicles but it does not localize to the edge of lamellipodia. The LifeAct-mScarlet-I serves as a marker for F-actin. Representative movie shown from three independent biological repeats. Cells were imaged every 10 seconds for the indicated times in minutes and seconds by wide-field time-lapse video microscopy using an IX 81 microscope (Olympus). Scale bar: 20 mm.

**Supplemental movie 6: EGFP-NHSL1-A1-WCA-DA localise to the edge of lamellipodia and vesicles**

This movie shows that EGFP-NHSL1-A1-WCA-DA expressed together with LifeAct-mScarlet-I in B16-F1 cells plated on laminin localises to the edge of lamellipodia and in addition to F-actin-rich “aster” structures from which vesicles appear to bud off in addition to its lamellipodia localisation. The LifeAct-mScarlet-I serves as a marker for F-actin. Representative movie shown from three independent biological repeats. Cells were imaged every 10 seconds for the indicated times in minutes and seconds by wide-field time-lapse video microscopy using an IX 81 microscope (Olympus). Scale bar: 20 mm.

## Notes

### Competing Interest Statement

The authors have declared no competing interest.

